# The mechanism of MinD stability modulation by MinE in Min protein dynamics

**DOI:** 10.1101/2021.07.30.454523

**Authors:** William C Carlquist, Eric N Cytrynbaum

## Abstract

The patterns formed both *in vivo* and *in vitro* by the Min protein system have attracted much interest because of the complexity of their dynamic interactions given the apparent simplicity of the component parts. Despite both the experimental and theoretical attention paid to this system, the details of the biochemical interactions of MinD and MinE, the proteins responsible for the patterning, are still unclear. For example, no model consistent with the known biochemistry has yet accounted for the observed dual role of MinE in the membrane stability of MinD. Until now, a statistical comparison of models to the time course of Min protein concentrations on the membrane has not been carried out. Such an approach is a powerful way to test existing and novel models that are difficult to test using a purely experimental approach. Here, we extract time series from previously published fluorescence microscopy time lapse images of *in vitro* experiments and fit two previously described and one novel mathematical model to the data. We find that the novel model, which we call the Asymmetric Activation with Bridged Stability Model, fits the time-course data best. It is also consistent with known biochemistry and explains the dual MinE role via MinE-dependent membrane stability that transitions under the influence of rising MinE to membrane instability with positive feedback. Our results reveal a more complex network of interactions between MinD and MinE underlying Min-system dynamics than previously considered.

---

The Min system of *Escherichia coli* is one of the simplest known biological systems that demonstrates diverse complex spatiotemporal behavior. It consists of three proteins, MinC, MinD, and MinE, which together dynamically regulate the position of the cell division site. Interactions of MinD and MinE on the cellular membrane drive coupled oscillations of the three Min proteins from cell pole to cell pole [39, 25, 19, 20, 24, 30], while MinC locally inhibits the formation of the Z-ring, the contractile ring that divides the cell in two [9, 3, 23, 53, 20, 38]. The character of Min-protein oscillations differs with cell length and shape. In rod-shaped cells, coupled densities of MinD and MinE stochastically switch from cell pole to cell pole in short cells [12], oscillate regularly from cell pole to cell pole in mid-sized cells [39, 14, 12], and oscillate regularly from cell pole to midcell in long mutant cells [39, 14]. In round mutant cells, densities of MinD and MinE oscillate antipodally [7, 45], and in branched mutant cells, densities of MinD and MinE oscillate from branch to branch to branch [48]. MinD and MinE also form dynamic patterns *in vitro*. In artificial, rod-shaped, membrane-clad compartments, densities of MinD and MinE exhibit oscillatory behaviors like in cells [55]. Perhaps most strikingly, on supported lipid bilayers, densities of MinD and MinE undergo spatially uniform oscillations in time [28] and form into traveling waves [32, 28, 31, 50, 49], spiral waves [32, 28, 31, 50, 49], dynamic amoeba-like shapes [28, 49], snake-like projections [28], mushroom-like shapes [49], and bursts [49].

Biochemistry, crystallography, and mutant studies have together elucidated many details of the protein structures and reactions that underly the emergent macroscopic behavior of the MinDE system. They have found that cytosolic MinD monomers bind to ATP and form dimers [24, 54, 52], which bind to a phospholipid membrane [18, 24, 22, 30, 54, 32] and cooperatively recruit other cytosolic MinD dimers to bind to the membrane [30]. Additionally, cytosolic MinE dimers bind to MinD dimers on the membrane [18, 31], bind to the membrane [36], and stimulate ATPase activity in the bound MinD dimers, causing the MinD dimers to separate and dissociate from the membrane [21, 18, 24, 30] while the MinE dimers transiently remain bound to the membrane [17, 46, 49] before returning to the cytosol. This characterization of MinDE kinetics is depicted in the top panel of Fig. 1 and referred to here as the Asymmetric Activation Model (AAM).

**Figure 1:**
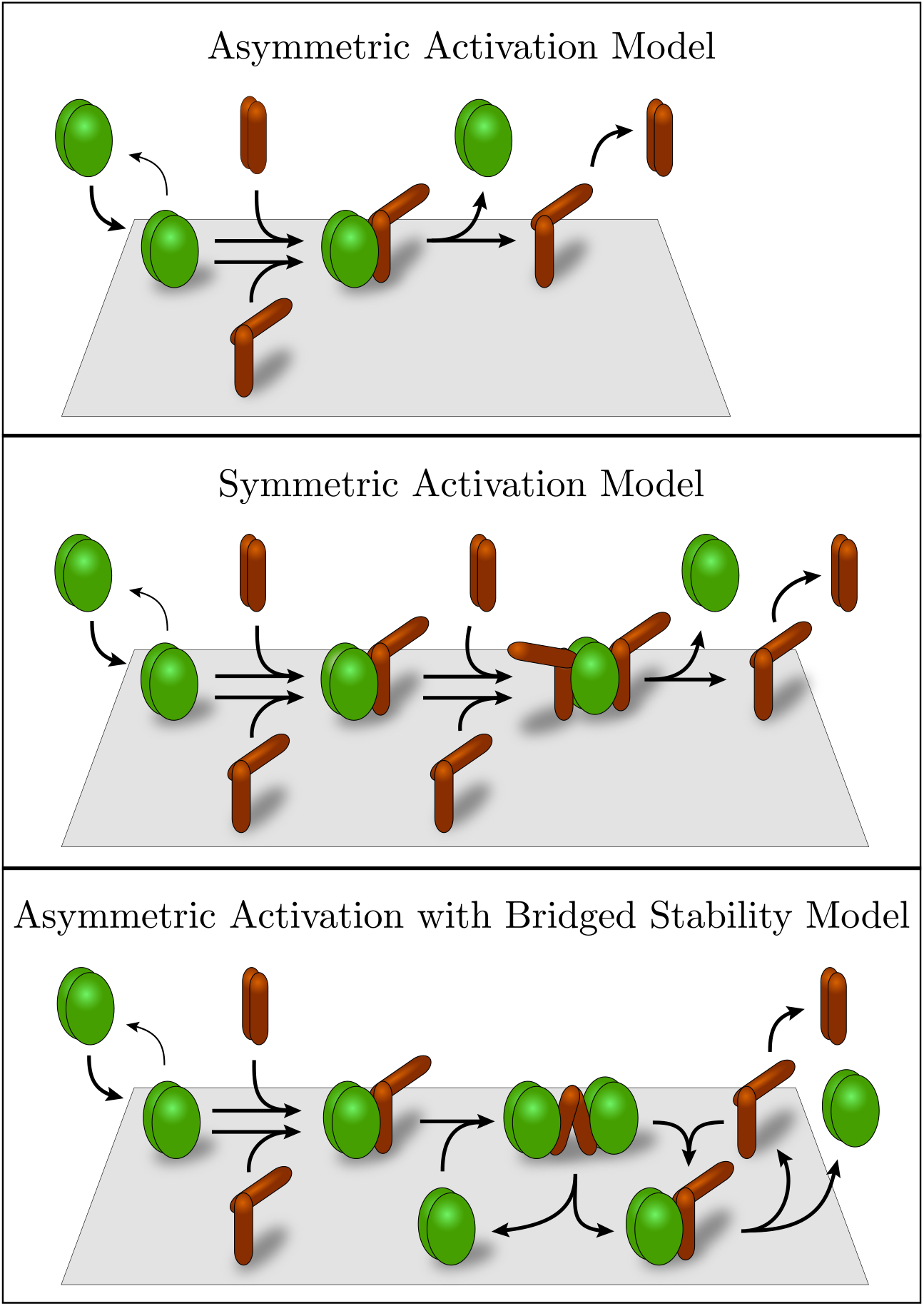
Cartoons of Min-system models considered - only conceptually important reactions are shown. Top panel: The AAM and CAAM - MinD dimers bind to the membrane and recruit MinE dimers which then induce ATP-hydrolysis in MinD dimers, causing MinD dimers to dissociate from the membrane. Middle panel: The Symmetric Activation Model. With a single MinE attached, ATP-hydrolysis in MinD dimers is not induced and MinD dimers are stabilized on the membrane. With two MinEs attached, hydrolysis is induced. Bottom panel: The Asymmetric Activation with Bridged Stability Model. This model includes the CAAM at its core but adds the possibility of a second MinD dimer binding to MinE, stabilizing both MinD dimers on the membrane. We hypothesize that the strain imposed on a MinE dimer from bridging two MinD dimers (note the changed conformation of MinE in the **ded** complex) alters MinE’s interaction with MinD, transiently disrupting its ability to induce ATP hydrolysis in either MinD dimer. Reaction details for the four models are given in Fig. 6

In bridging the gap between protein interactions and emergent patterning, mathematical modeling of the Min system has a long and rich history. Most mathematical models of MinDE dynamics are based on a subset of the reactions described above, and generally their aim has been to recapitulate behaviors of the Min system *in vivo*. They have successfully demonstrated oscillatory behaviors particular to short cells [12, 4], mid-sized cells [16, 34, 29, 43, 51], long cells [34, 26, 33, 47, 4, 51], dividing cells [47, 44, 10, 51], aberrant cellular geometries [27, 11, 13, 4, 48, 41], and MinE mutants [8, 2]. Several mathematical models have recapitulated behaviors of the Min system *in vitro*, including traveling waves [32, 37] and spiral waves [32, 4] on supported lipid bilayers and patterning on geometrically confined membranes [42] and micropatterned substrates [15].

Various quantitative experimental measurements have been used to validate different mathematical models of the Min system: pole-to-pole oscillation period in vivo [33, 47, 8, 5, 2, 13, 4, 51], distributions of residence times during stochastic pole-to-pole switching [12, 4] and regular pole-to-pole oscillations [12] *in vivo*, and traveling wave velocity and wavelength *in vitro* [32]. However, no model has been quantitatively compared to time-course data. A quantitative comparison of how well models fit time-course data is the next logical step in model selection and validation.

In Ivanov and Mizuuchi’s *in vitro* experiments [28], buffer was flowed atop a supported lipid bilayer to ensure spatially uniform concentrations of reaction components in the buffer. On the supported lipid bilayer, densities of MinD and MinE oscillated nearly homogeneously in space before forming into traveling waves. Under spatially uniform conditions, systems of partial differential equations (PDEs) that model how Min protein concentrations evolve in space and time reduce to systems of ordinary differential equations (ODEs) that describe just the local reactions. This set-up greatly simplifies the modelling. We find regions of space in which the oscillation data from Ivanov and Mizuuchi’s experiments have the least amount of spatial variation and extract one period of the spatially near-homogeneous oscillation time-course data from it, hereafter referred to as *oscillation data*. Fitting ODE models to the oscillation data allows us to compare different reaction-mechanism models without complications from spatial effects but still giving insight into patterning. The oscillation data are shown as dots in the top panel of Fig. 2, and details of data extraction are discussed in the Methods.

**Figure 2:**
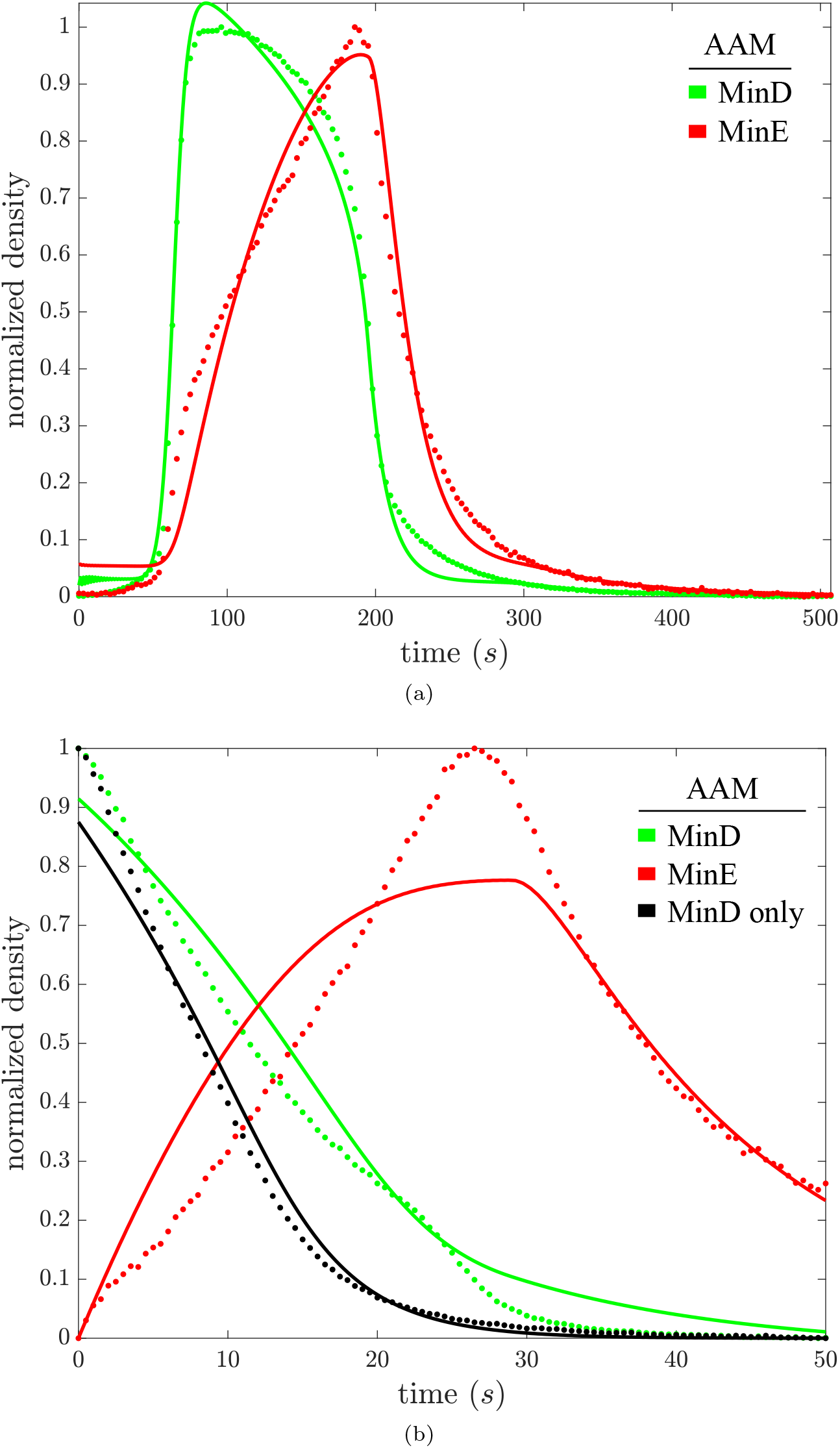
For the Asymmetric Activation Model (AAM), the fit to the oscillation data is shown in (a), and the fit to MinD dissociation data is shown in (b). Fits are solid, data are dotted. The data has been scaled and shifted so as to span 0 to 1, and the model has been modified with the same transformation in this and the next figure to ensure fits to both MinD and MinE time courses are clearly visible.

Vecchiarelli *et al*. recently showed that, in addition to acting as an inhibitor of MinD membrane binding, at times MinE can stabilize MinD on the membrane [49]. In *in vitro* experiments similar to those of Ivanov and Mizuuchi, they flowed buffer with and without MinE atop a layer of MinD bound to a supported lipid bilayer. Initially, MinD dissociated more slowly from the membrane with MinE in the flowed buffer than it did without MinE in the flowed buffer. Later, the concentration of membrane-bound MinE increased and MinD dissociated more rapidly from the membrane than it did without MinE in the flowed buffer. Fitting ODE models to Vecchiarelli *et al*.’s MinD dissociation data with and without MinE in the flowed buffer allows us to compare how well different reaction-mechanism models recapitulate MinE’s dual role in MinD membrane binding. The MinD dissociation data with and without MinE in the flowed buffer, hereafter referred to as *MinD dissociation data*, are shown as dots in the bottom panel of Fig. 2. We refer to the oscillation data and the MinD dissociation data together as the *time-course data*.

The mechanism underlying MinE’s dual role in MinD membrane binding is unclear, no mathematical model has accounted for it, and its biological implications remain unknown. The only current hypothesis proposes that one bound MinE dimer anchors a MinD dimer to the membrane, whereas two bound MinE dimers stimulate ATPase activity in a MinD dimer, causing it to separate and dissociate from the membrane [49]. We formulate this mechanism into a mathematical model that we call the Symmetric Activation Model (SAM). Mutational studies have shown, however, that a single MinE dimer bound to a MinD dimer is sufficient to stimulate ATPase activity in the MinD dimer [35]. Based on this finding, we propose a new model, the Asymmetric Activation with Bridged Stability Model (AABSM), that accounts for MinE’s dual role in MinD membrane binding and requires only one bound MinE dimer to stimulate ATPase activity in a MinD dimer. Both these models are shown in a simplified form in Fig. 1 with details given in the Methods.

We fit the SAM and the AABSM to the oscillation data and the MinD dissociation data using the Homotopy-Minimization Method for parameter estimation in differential equations [6]. For comparison, we also fit a mathematical version of the AAM to the time-course data as well as a Comprehensive Asymmetric Activation Model (CAAM) and a generic excitability model (the FitzHugh-Nagumo model). Examining how models fit the time-course data allows us to distinguish between biochemical assumptions in the various models. Our model fitting also provides us with time courses for hidden protein states that would be difficult or impossible to measure experimentally. Ultimately, our analysis supports the AABSM and suggests that, through a more complex web of interactions, activations, and inhibitions between MinD and MinE than previously considered, Min-system pattern formation is driven by a MinE-dependent reinforced stability of MinD on the membrane followed by a MinE-dictated switch to membrane instability with positive feedback.

## Results

### Fitting The Asymmetric Activation Model

Amongst previously published mathematical models, the model of Bonny *et al.* [4] (henceforth the Bonny Model) demonstrates the most diverse array of behaviors that are qualitatively similar to experimental observations of the Min system *in vivo* and *in vitro*, including stochastic pole-to-pole switching in short cells, regular pole-to-pole oscillations in mid-sized cells, oscillation splitting in growing cells, regular pole-to-midcell oscillations in long cells, end-to-end oscillations in thick cells, and spiral waves on a supported lipid bilayer. In the Bonny Model, bulk MinD dimers bind to the membrane (**D → d**), and membrane-bound MinD dimers cooperatively recruit more bulk MinD dimers to bind to the membrane 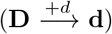 or bind to bulk MinE dimers forming membrane-bound MinD-MinE complexes (**d,E → de**). Then, MinE dimers in the **de** complex stimulate ATPase activity in bound MinD dimers, causing the MinD dimers to dissociate from the membrane while the MinE dimers either return to the bulk (**de → D,E**) or temporarily remain on the membrane (**de → D,e**). Finally, membrane-bound MinE dimers either dissociate from the membrane (**e → E**) or bind to membrane-bound MinD dimers forming membrane-bound MinD-MinE complexes (**d,e → de**). Beyond the aforementioned reactions, bulk species diffuse in the bulk and membrane-bound species diffuse on the membrane in the Bonny Model. Given the Bonny Model’s ability to qualitatively recapitulate Min-system patterning, we wanted to see how well it could describe the time-course data.

We modified the Bonny Model to be consistent with experimental conditions and outcomes underlying the time-course data to get the AAM, so named to emphasize that a single MinE dimer is sufficient to activate hydrolysis in the MinD dimer. Details of the modifications are discussed in the Methods. A cartoon showing some of the defining characteristics of the AAM is shown in the top panel of Fig. 1. The full set of reactions in the AAM are depicted in Fig. 6 and the AAM is written as a system of ODEs in Eq. S2.

Fitting the AAM to the time-course data, as described in the Material and Methods, we found reasonably good qualitative agreement, but certain details differed as can be seen in Fig. 2. For the oscillation data, after peaking too high, the fit to the MinD density drops too quickly at first, has a similar maximal rate of decrease compared to the time-course data, and then drops too quickly again in the tail of the pulse. Additionally, the AAM also fails to recapitulate the near linear rise in MinE density from roughly 70 s until 190 s. As with MinD, the final decay of MinE back to steady state is also too rapid. For the MinD dissociation data with MinE in the flowed buffer, as was expected given that the AAM treats MinE solely as an inhibitor of MinD membrane binding, the AAM fails to recapitulate the knee in MinD density around 25 s, when MinE shifts from stabilizing to destabilizing MinD on the membrane. Additionally, the AAM fails to recapitulate the increasing rate of net MinE attachment from roughly 5 s until 20 s. Instead, the MinE density rises quickly at first and then tapers off, seemingly, as the amount of MinD available for it to bind to on the membrane drops off. Furthermore, we note that although the AAM appears to qualitatively describe the time-course data fairly well, a variant of the FitzHugh-Nagumo Model (FHNM), a simplified model of neuron firing, qualitatively describes the time-course data almost as well as the AAM, as can be seen in Fig. S4. As such, we caution that what appears to be a reasonably good qualitative agreement with data does not necessarily qualify as a good biochemical model. In Table 1, we show the relative *χ*^2^ statistic and Akaike Information Criterion score (AIC) for all models considered. These quantities clarify that neither the FHNM nor the AAM do particularly well in explaining the time-course data, but that the AAM does better than the FHNM.

**Table 1:** Model Comparison. *χ*^2^ is the the weighted sum of squared residuals, AIC is the Akaike Information Criterion score, 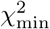 is the minimum *χ*^2^ value amongst the models, AIC_min_ is the minimum AIC value amongst the models, and n is the number of fitting parameters. The AIC is an estimate of the information lost by using the model to represent the data and accounts for the fitting benefit of having more parameters; it decreases as the fit to data improves and increases with the number of parameters in the model. As a rule of thumb, a difference in AIC of more than 10 between two models is strong evidence in favor of the model with the lower AIC score [1]. By both *χ*^2^ and AIC measures, the AABSM clearly describes both sets of time-course data best.

|  | Oscillation Data |  |  | MinD Dissociation Data |  |  |
| --- | --- | --- | --- | --- | --- | --- |
| | $\chi^2/\chi_{\min}^2$ | AIC – AIC<sub>min</sub> | $n$ | $\chi^2/\chi_{\min}^2$ | AIC – AIC<sub>min</sub> | $n$ |
| FHNM | 90 | 1500 | 9 | 260 | 1600 | 10 |
| AAM | 52 | 1300 | 14 | 150 | 1500 | 13 |
| CAAM | 14 | 880 | 18 | 15 | 800 | 16 |
| SAM | 1.8 | 200 | 29 | 3.2 | 360 | 26 |
| AABSM | 1 | 0 | 28 | 1 | 0 | 25 |

### Fitting The Comprehensive Asymmetric Activation Model

Based on experimental observations since the publication of the Bonny Model, we extended the AAM to include additional reactions (see the Supplemental Material for details), then fit the Comprehensive Asymmetric Activation Model (CAAM) to the time-course data to see if the inclusion of the new reactions markedly improves data fitting. A cartoon showing some of the defining characteristics of the CAAM is shown in the top panel of Fig. 1, the full set of reactions in the CAAM are depicted in Fig. 6, and the CAAM is written as a system of ODEs in Eq. S3.

We found that the CAAM fits the time-course data substantially better than the AAM, as can qualitatively be seen in Fig. S2 and is quantitatively shown by *χ*^2^ statistics and AIC scores in Table 1.

The reaction 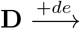 allows recruitment of MinD to the membrane throughout the pulse, and its inclusion in the CAAM appears to have alleviated a peak overshoot followed by an accelerated drop-off of MinD in the fit to the oscillation data. The inclusion of facilitated MinE recruitment (reactions 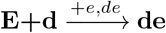) in the CAAM appears to have allowed for a better match to the observed constant rate of net MinE-membrane attachment between 70 s and 190 s in the oscillation data and the increasing rate of net MinE-membrane attachment from roughly 5 s until 20 s in the MinD dissociation data with MinE in the flowed buffer, presumably by increasing facilitation even as the bindable amount of MinD on the membrane decreases. Ultimately, the CAAM appears to fit the oscillation data fairly well. However, like the AAM, the CAAM is unable to demonstrate MinE’s dual role as both a stabilizer and a destabilizer of MinD membrane binding.

### Fitting The Symmetric Activation Model

To account for MinE’s dual role as both a stabilizer and a destabilizer of MinD membrane binding, Vecchiarelli *et al.* hypothesized that the binding of a single MinE dimer to a membrane-bound MinD dimer is not sufficient to stimulate ATPase activity in the MinD dimer [49]. Rather, they proposed that the binding of a single MinE dimer to a membrane-bound MinD dimer stabilizes the MinD dimer on the membrane through MinE’s interaction with the membrane, and the subsequent binding of a second MinE dimer to the membrane-bound MinD dimer stimulates ATPase activity in the MinD dimer, causing it to dissociate from the membrane. A simplified version of this biochemical arrangement was also proposed previously in a modeling study with different motivation before it was known that MinE played multiple roles in MinD’s interaction with the membrane [37]. No other hypothesis has been proposed to explain MinE’s dual role in MinD-membrane binding.

We formulated Vecchiarelli *et al.*’s hypothesis in a mathematical model that we call the Symmetric Activation Model (SAM) because it requires two MinE dimers, one attached to each MinD-dimer subunit, to induce removal of a MinD dimer from the membrane. In doing so, we included a wide range of possible reactions in the SAM because we had no experimental basis for the full set of reactions that should be included in it. Details are given in the Methods. A cartoon showing some of the defining characteristics of the SAM is shown in the middle panel of Fig. 1, the full set of reactions in the SAM are depicted in Fig. 6, and the SAM is written as a system of ODEs in Eq. S4.

We found that the optimal data-fitting parameter choice for the SAM provides a better fit to both sets of time course data than the CAAM (see Fig. S3 and Table 1). Most notably, for the MinD dissociation data with MinE in the flowed buffer, the SAM partially recapitulates the knee in MinD density around 25 s, when MinE shifts from stabilizing to destabilizing MinD-membrane binding, whereas the CAAM does not. However, the SAM demonstrates a less dynamic shift than is visible in the data.

### Fitting The Asymmetric Activation with Bridged Stability Model

Contradicting the underlying assumption of the SAM, experiments have shown that a single MinE dimer bound to one subunit of a MinD dimer stimulates ATPase activity in both subunits of the MinD dimer [35], supporting the theory that MinE dimers stimulate ATPase activity in MinD dimers asymmetrically rather than symmetrically. There is no direct evidence for the structure of a membrane-stable MinD-MinE complex, but one crystal structure shows a MinE dimer bridging two MinD dimers. In this structure, one MinD dimer is rotated 90° with respect to the other MinD dimer [36], which has led to the structure being considered an experimental artifact rather than a biologically relevant state. Despite this, we propose that a MinE dimer bridging two MinD dimers (**ded**) may be the membrane-stable MinD-MinE complex. We hypothesize that the strain associated with the 90° rotation required to ensure both MinD dimers can bind to the membrane inhibits ATPase activity in both MinD dimers to a level below the activity in a **de** complex, by perturbing both MinD-MinE interfaces from that in the **de** complex. Furthermore, we assume that the additional membrane anchors in the **ded** complex stabilize the complex as compared to the **d**-dimer. This latter assumption provides an explanation for the slower observed detachment of MinD in the presence of MinE as compared to in its absence. These assumptions guided our formulation of the AABSM.

Like in the SAM, we included a wide range of possible reactions in the AABSM. A cartoon showing some of its defining characteristics is shown in the bottom panel of Fig. 1. The full set of AABSM reactions are depicted in Fig. 6, and the model is written as a system of ODEs in Eq. S5.

We found that the AABSM fits both time-course data sets better than the SAM (see Fig. 3 and Table 1) and could describe all features of the time-course data, including the knee in MinD density at 25 s in the MinD dissociation data with MinE in the flowed buffer when MinE shifts from MinD-stabilizing to MinD-destabilizing, a feature that was not well captured by any of the other three models.

**Figure 3:**
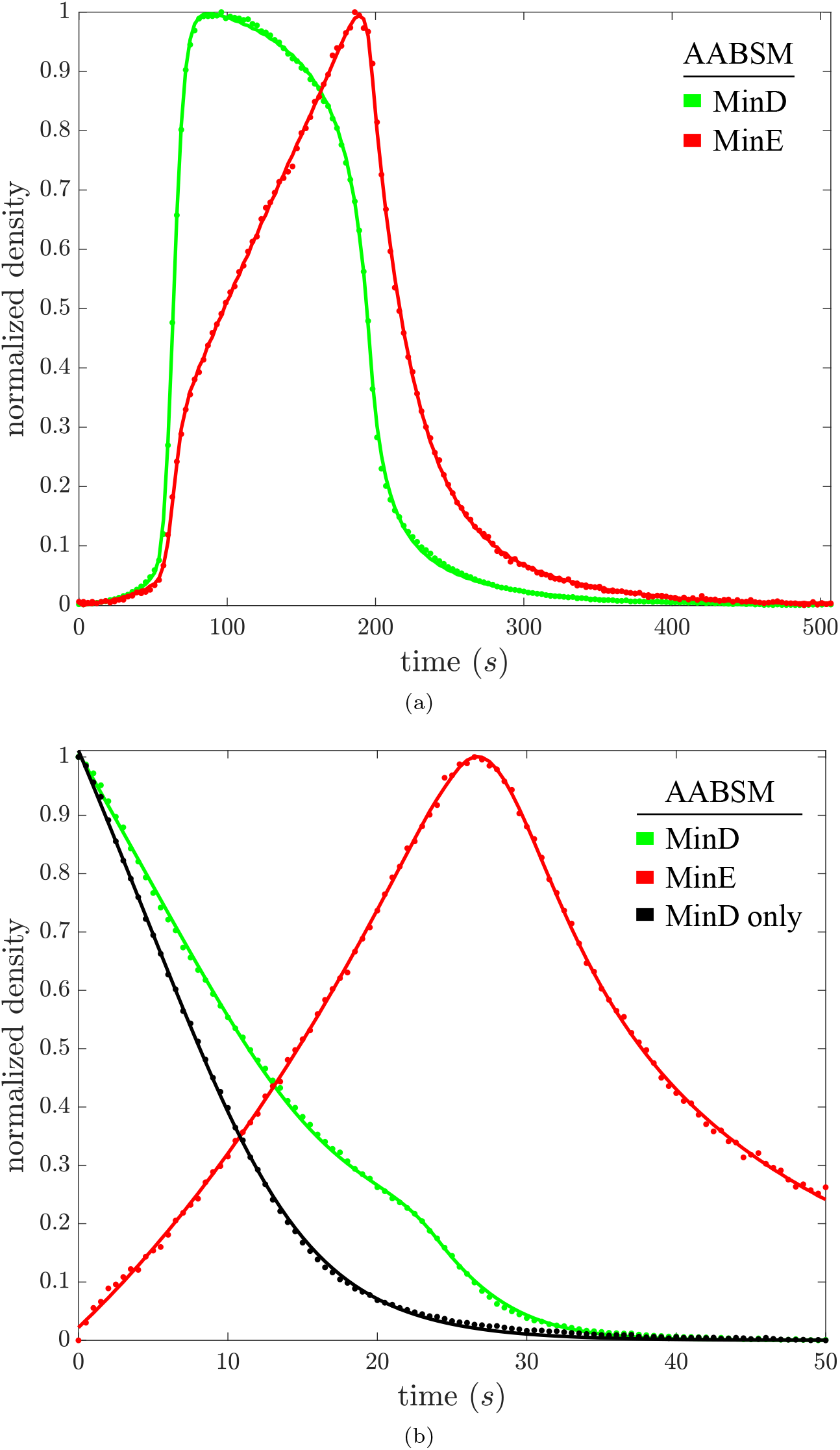
For the Asymmetric Activation with Bridged Stability Model (AABSM), the fit to the oscillation data is shown in (a), and the fit to MinD dissociation data is shown in (b). Data is shown as dots with colors for MinD and MinE matching the model colors.

State values from the fits of the AABSM to the time-course data are shown in Fig. 4. When concentrations of membrane-bound MinD dimers (green curve) are high, MinE is predominantly bound by two MinD dimers (blue curve) and MinD is stable on the membrane. As the concentration of membrane-bound MinD dimers decreases, the predominant form of MinE is bound to a single MinD dimer (red curve) and hence MinD is unstable on the membrane. Additionally, only when concentrations of membrane-bound MinD dimers become negligible (at about 200 s in the oscillation data and about 24 s in the MinD dissociation data with MinE in the flowed buffer) do the concentrations of membrane-bound MinE dimers (black curve) become non-negligible and persist. This progression of membrane-bound states is described in more detail in the Discussion.

**Figure 4:**
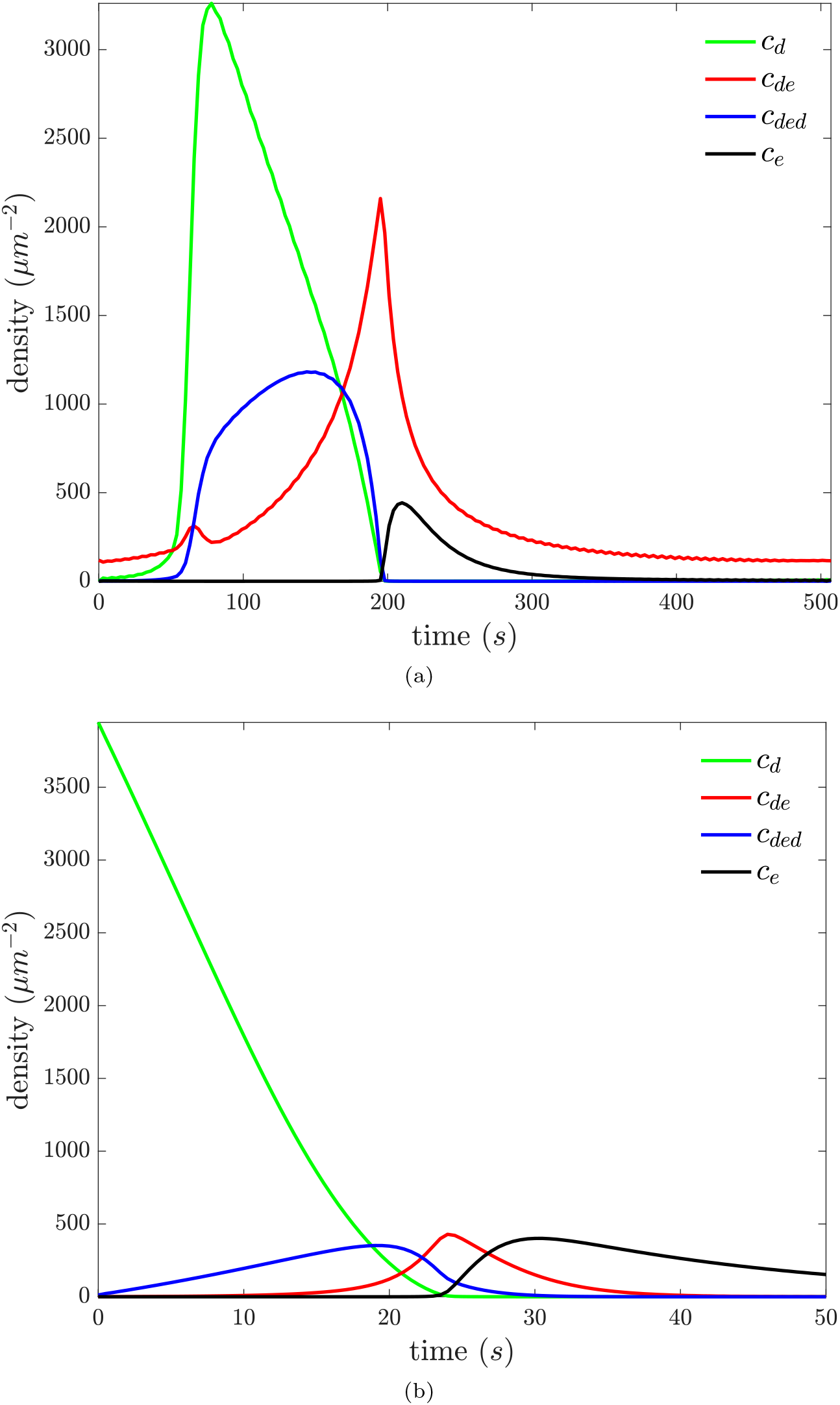
For the AABSM, state values from the fit to the oscillation data are shown in (a), and state values from the fit to MinD dissociation data with MinE in the flowed buffer are shown in (b). *c*_*d*_ is the concentration of membrane-bound MinD dimers (**d**), the membrane-semistable MinD state; *c*_*de*_ is the concentration of MinE dimers bound to one MinD dimer (**de**), the membrane-unstable MinD state; *c*_*ded*_ is the concentration of MinE dimers bridging two MinD dimers (**ded**), the membrane-stable MinD state; and *c*_*e*_ is the concentration of membrane-bound MinE dimers (**e**). We note that the concentration of membrane-stable MinD dimers is twice the value of *c*_*ded*_. One important feature seen in both plots is the progression from green to blue to red to black, which emphasizes the ordered dynamics.

### Removed-Reaction Fits of The AABSM

Without direct experimental evidence for the set of reactions that should comprise the AABSM, we included a large range of possible reactions in it. To clarify which reactions are important for describing the time-course data, we individually removed each non-necessary reaction from the AABSM and determined how well the resulting model could fit the time-course data. From a large decrease in the quality of fit we infer that the removed reaction is important, whereas a negligible decrease suggests that the removed reaction is either unimportant in describing the time-course data or its effect can be compensated for by altering other parameters in the model. See Table 2 for the influence of each removed reaction on the AABSM’s *χ*^2^ value and AIC score.

**Table 2:** A single-reaction knock-out study for the AABSM. We removed each listed reaction in turn from the AABSM and re-optimized the parameter values to get the best fit to the time-course data. For each resulting model, *χ*^2^ is the the weighted sum of squared residuals, and AIC is the Akaike Information Criterion. 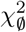 and AIC_Ø_ are the weighted sum of squared residuals and the Akaike Information Criterion for the AABSM with all reactions included. Greater values of 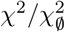 and AIC – AIC_Ø_ correspond to a larger decrease in the quality of the fit. The reactions are ordered b;y their AIC difference from the complete AABSM for the oscillation data with related reactions grouped together. See the caption of Table 1 for a discussion of AIC scale.

| Removed reaction | Oscillation Data |  | MinD Dissociation Data |  |
| --- | --- | --- | --- | --- |
| | $\chi^2/\chi_\emptyset^2$ | $\text{AIC} - \text{AIC}_\emptyset$ | $\chi^2/\chi_\emptyset^2$ | $\text{AIC} - \text{AIC}_\emptyset$ |
| **ded+e → de+de** | 4.8 | 510 | 1.0 | -1.5 |
| **de → D+E** | 4.2 | 480 | 1.5 | 130 |
| **de → d+e** | 1.0 | 4.6 | 1.0 | -2.0 |
| **de → D+e** | 1.0 | -6.0 | 1.0 | 8.0 |
| **d+de → ded** | 4.0 | 460 | 1.2 | 64 |
| **D $\xrightarrow{+de}$ d** | 3.5 | 410 | NA | NA |
| **D $\xrightarrow{+ded}$ d** | 2.0 | 240 | NA | NA |
| **D $\xrightarrow{+d}$ d** | 1.8 | 200 | NA | NA |
| **E+d $\xrightarrow{+e}$ de** | 2.7 | 330 | 1.0 | 9.8 |
| **E+d $\xrightarrow{+ded}$ de** | 1.0 | 3.1 | 1.0 | -0.21 |
| **E+d $\xrightarrow{+de}$ de** | 1.0 | 0.23 | 1.0 | -2.2 |
| **de+de → ded+e** | 1.5 | 140 | 1.1 | 33 |
| **d+e → de** | 1.3 | 75 | 1.1 | 32 |
| **E+ded $\xrightarrow{+e}$ de+de** | 1.2 | 49 | 1.1 | 15 |
| **E+ded $\xrightarrow{+ded}$ de+de** | 1.2 | 48 | 1.1 | 18 |
| **E+ded → de+de** | 1.0 | -1.2 | 1.0 | 1.7 |
| **E+ded $\xrightarrow{+de}$ de+de** | 1.0 | -0.17 | 1.1 | 23 |
| **ded → d+de** | 1.0 | -1.3 | 1.1 | 19 |

As seen in Table 2, the reactions **ded+e** → **de+de** and 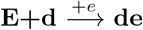 are critical for the AABSM to recapitulate the oscillation data well. Although the table indicates other important reactions, we highlight these two because they suggest previously unknown functions of MinE dimers and for their role in the fundamental mechanism elaborated in the discussion section.

Additionally, we note that overall the AABSM is far more sensitive to reaction removal with the oscillation data than with the MinD dissociation data.

## Discussion

Over the last twelve years, the success of reconstituting the Min protein interactions on supported membranes has allowed for new insights into the behavior of the system [32, 31, 28, 49]. Some of these experimental studies have been accompanied by mathematical modelling work, but none of these modelling studies nor others have attempted validation against time courses of protein concentration, instead focusing on qualitative features (e.g. pattern type, bifurcations, and transitions from stochastic to deterministic dynamics) and macroscopic quantification (e.g. oscillation period, wavelength, and wave velocity) [32, 12, 4, 37]. We formulated a biochemically novel model and used model selection, taking advantage of recent high quality quantitative time-course data [28, 49], to evaluate it against other recent models. Our model performed significantly better than the others, providing excellent fits. It is also consistent with other recent biochemical observations, sheds lights on the functional roles of MinE domains, and provides insight into factors that control the underlying dynamics of Min-system patterning, as discussed below.

### A free-MinD controlled stability-switching dynamic underlying patterning

Throughout the description below, our claims are based on a combination of the model comparisons, the time-course of state values in the AABSM shown in Fig. 4, and the removed-reaction results in Table 2.

The biochemistry of the AABSM has the membrane stability of three forms of the MinD-membrane bond at its core. The least membrane-stable form is the MinD-MinE (**de**) complex. The most stable is the MinD-MinE-MinD (**ded**) complex. Intermediate to these two is the MinD dimer on its own (**d**).

The dynamics of the model start with a wave of MinD dimer attachment to the membrane (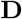 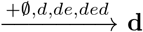 [see reactions in Table 2], green upstroke [see colored curves in Fig. 4a]). This precedes and coincides with MinE recruitment and **de** complex formation (**d+E → de**, the first small upstroke in red). While the MinD dimer concentration is high, **de** complexes are quickly sequestered into the stable **ded** complex (**d+de → ded**, the drop in red accompanied by the upstroke and subsequent plateau in blue). This **ded**-complex plateau is much like a short-lived quasi-stable state seen in many excitable media. During this quasi-stable period, the flow of the **ded** complex into the **de** state, which is important later in the cycle, is suppressed by the initial sequestration of MinE in the **ded** state (**2d+e → d+de → ded**). This MinE-reinforced stability is illustrated in the top panel of Fig. 5.

**Figure 5:**
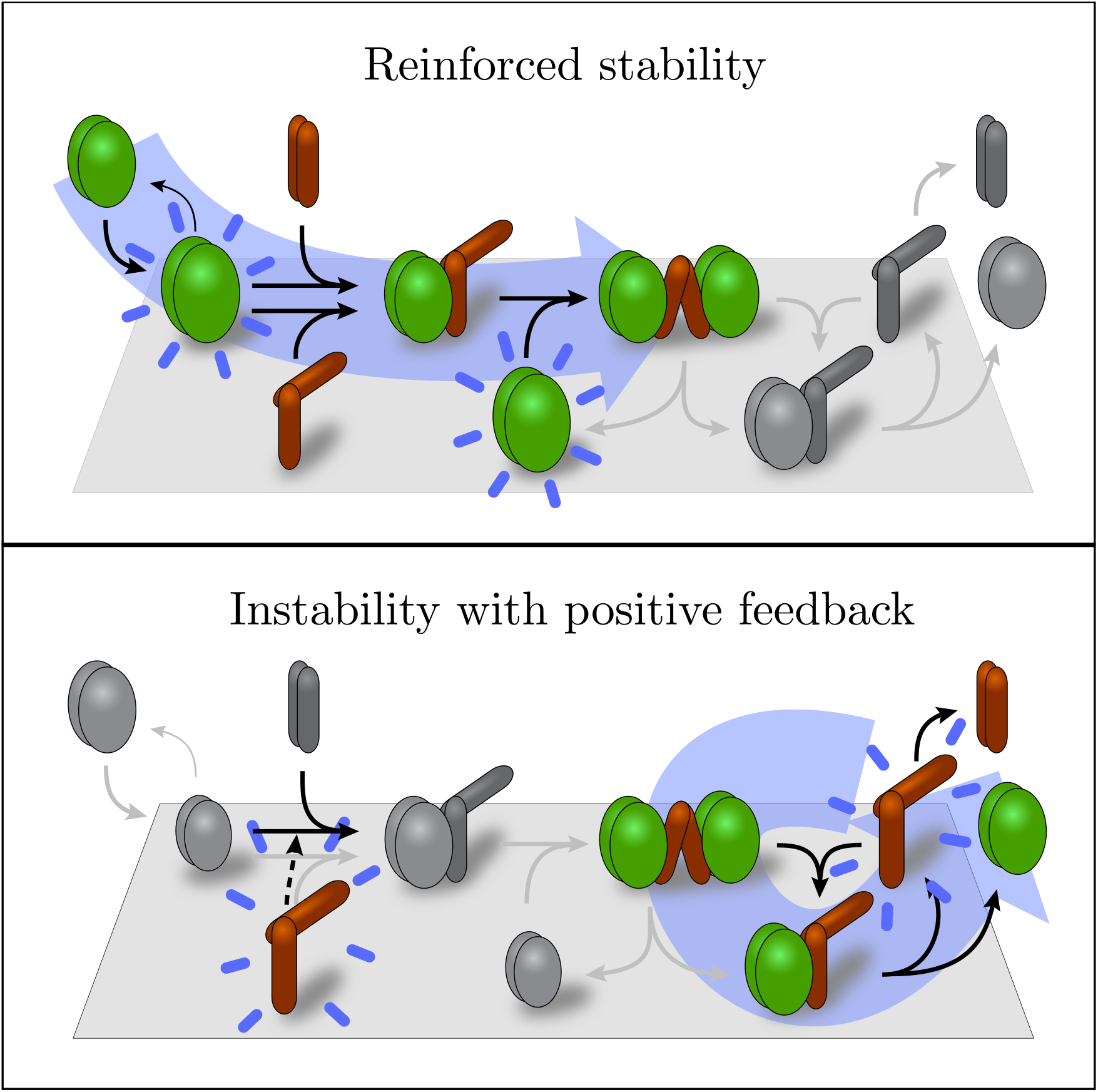
The membrane cycling of the Min proteins can be separated into two phases. In the first phase (reinforced stability), MinD attaches in greater numbers than MinE, and all forms of MinE are shuttled into the membrane-stable **ded** form (blue arrow in the top panel). The stability is reinforced by the resultant depletion of **e**, preventing the destabilizing feedback that happens in the second phase (instability with positive feedback). In the second phase, membrane saturation prevents further MinD attachment, allowing MinE recruitment to catch up. Two forms of positive feedback accelerate the removal of MinD - **e** breaks the membrane-stable **ded** into two membrane-unstable **de**’s which subsequently leave more **e** on the membrane as MinD is removed (blue arrow in bottom panel), and **e** recruits additional MinE from the bulk to bind to **d** forming **de** (dashed line) which, again, thereafter deposits more **e** on the membrane as MinD dissociates. In both panels, the state that drives the phase’s behavior is highlighted with a blue aura, and the relative concentration of **d**, which dictates overall stability, is shown by size.

The next important transition occurs when new **d** recruitment is prevented by membrane saturation and MinE recruitment catches up. This change in the overall MinD-to-MinE ratio tips the balance from sequestration (**d+de → ded**) to MinD removal (**ded → d+de → d+D+e**), leaving some of the MinE dimer behind. The additional **e** from the removal step accelerates the previously suppressed desequestration (**ded+e** → **de+de**) to a flood (precipitous drop in blue as red spikes). This combined with the reaction 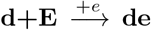 pushes the membrane-stable MinD states into the **de** state, with subsequent MinD removal. The flood of **de** complexes is only possible once the MinD dimer supply is low and hydrolysis provides a supply of membrane-bound MinE. Thus the MinD-to-MinE ratio acts as a control parameter that dictates whether the membrane-stable **ded** complex or the membrane-unstable **de** complex is the dominant form of MinD. This positive feedback of MinE destabilizing MinD is illustrated in the bottom panel of Fig 5.

In contrast with previous models, here we have delved further into the unknown dynamics of the experimentally hidden states and used model selection to sort out the most likely progression through those states. The idea that MinD moves into a meta-stable state (**ded**), reinforced by inhibition of the backward reaction (sequestering), followed by destabilization under positive feedback is a refinement of the simple “D attaches, E removes” paradigm that makes strong falsifiable predictions about the underlying biochemistry.

### Biochemical observations supporting the AABSM over the SAM

Evidence to support the AABSM’s membrane-stability assumptions comes from a few sources. It has been long known that MinE can induce hydrolysis of ATP by the MinD dimer [18]. The AABSM adheres to the conventional modelling interpretation that it is the MinD-MinE (**de**) complex that is responsible for the induction of hydrolysis, given that a single MinE dimer bound to a MinD dimer is capable of inducing hydrolysis in both subunits of the MinD dimer [35]. This latter observation is at odds with the membrane-stability assumptions of the SAM which has the **de** complex as stable on the membrane.

The assumption of reduced hydrolysis in the MinD-MinE-MinD (**ded**) complex is more speculative but is nonetheless supported by experiments. Vecchiarelli *et al.* showed that the MinD ATPase rate as a function of MinE concentration is sigmoidal and that the deletion of the dimerization domain of MinE removes the sigmoidal nature of the curve [49]. The sigmoidal MinE dependence is consistent with both the SAM and the AABSM. The loss of that sigmoidal dependence with the deletion is what the AABSM would predict, but the SAM would require some additional assumptions about the role of the second MinE subunit to be consistent with the loss.

In the SAM and the AABSM, a high concentration of MinE under mass action would leave negligible MinD on the membrane in the unbound dimer form, **d**, and would push all MinD into the model’s respective membrane-unstable state, **ede** for the SAM or **de** for the AABSM. Apart from forming complexes with MinD on the membrane, some MinE may also be present in the **e**state in both models. As such, a high concentration of MinE should push the ratio of MinD to MinE on the membrane to be no greater than 1/2 for the SAM and no greater than 1 for the AABSM. In a Min-system assay using a non-hydrolyzable ATP analogue testing precisely this, the MinD-to-MinE ratio on bicelles was found to be 1 [24], more in line with the prediction of the AABSM than the SAM.

### MinE membrane-binding and dimerization

Our results shed light on the critical but unknown role of MinE-membrane binding. One interpretation is that MinE-membrane binding allows MinE to remain on the membrane more persistently, permitting a MinE dimer to stimulate ATPase activity in multiple MinD dimers before dissociating from the membrane [36, 35]. Our supported model and removed-reaction fits suggest additional roles via the membrane-bound MinE state (**e**) in the reactions 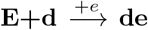 and **e+ded** → **2de**. On a supported-lipid-bilayer *in vitro*, the MinE-membrane-binding-deficient mutant MinE C1 [17] together with MinD formed a stationary structure in contrast to the traveling wave seen with wildtype MinE [31]. The traveling and stationary wave forms are similar except that the MinE C1 profile is rounded in comparison to the sharp peak in wildtype MinE [31]. Drawing on our findings, the lack of a sharp peak in the MinE C1 profile could follow from the inability of MinE C1 to recruit bulk MinE C1 to bind to membrane-bound MinD (the first reaction above), and the formation of a stable, stationary MinD/MinE C1 structure on the supported lipid bilayer could follow from the absence of the destabilizing second reaction which would be disabled by the MinE C1 mutant.

MinE dimerization is required for patterning both *in vivo* [40] and *in vitro* [49], but a role for MinE dimerization has not, to our knowledge, been postulated. In our supported model, MinE dimerization is critical for the sequestration of MinD in the stable **ded** state. Without it, a (strained) MinE dimer at the center of the complex would not be able to bridge two MinD dimers. Given that the **ded** state lies at the core of our supported model, its removal through the elimination of MinE dimerization would presumably distrupt patterning both *in vivo* and *in vitro*, although the details of exactly how it would be disturbed are not clear. More concretely, a knock-out of the **ded** state provides an explanation for the observed loss of MinD stabilization in MinD-dissociation experiments when a dimerization-deficient MinE mutant is substituted for wildtype MinE (compare [49] Fig. 6B first and third panels).

**Figure 6:**
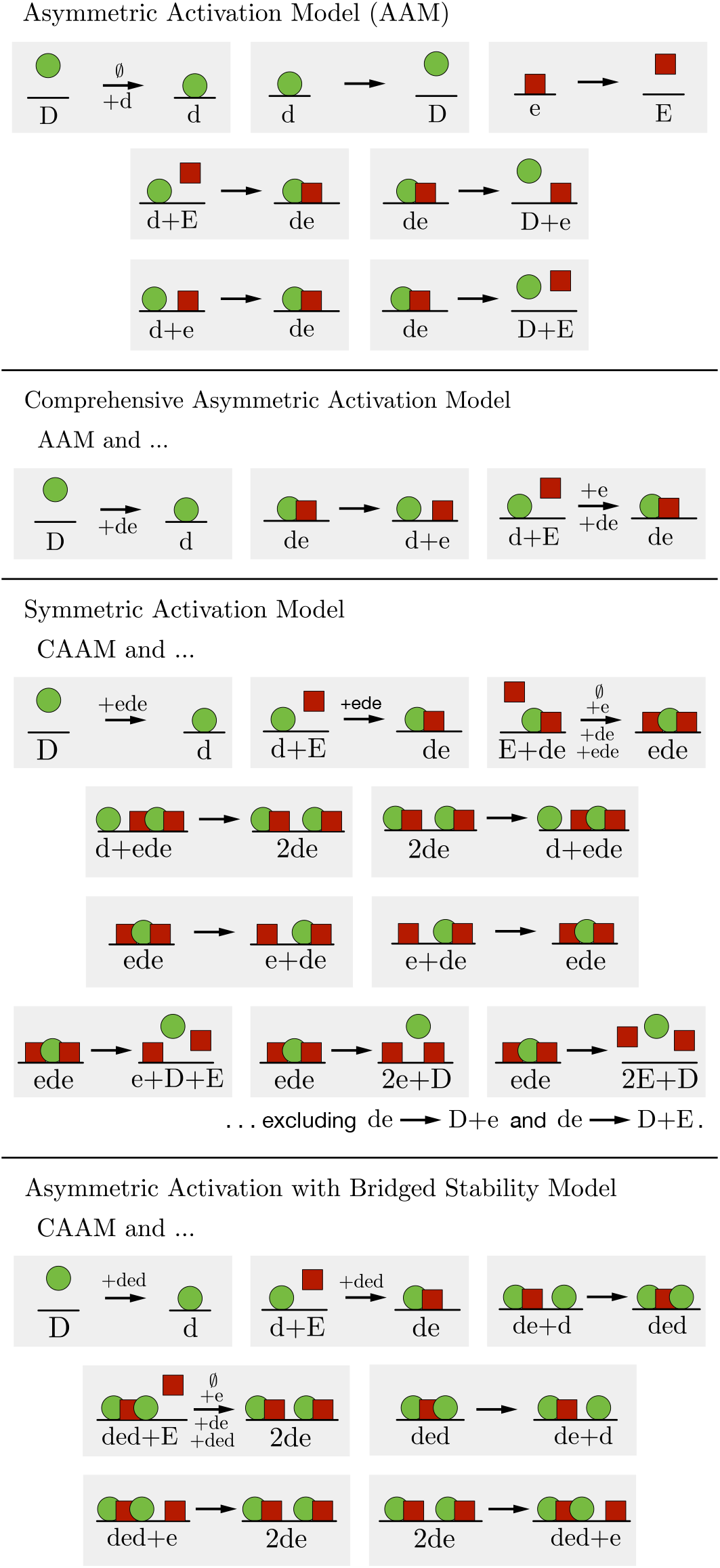
Reactions details for the four models. MinD is represented as a green circle, MinE as a red square, and the supported membrane as a black line. A reaction arrow annotated with a “+x” indicates that the reaction is facilitated by the density of protein (or protein complex) x. To indicate that the arrow also includes an unfacilitated basal rate, we annotate with *Ø*.

MinE membrane-binding and dimerization are integral to opposing processes in our supported model, the former required for destabilizing MinD and the latter required for stabilizing it. As such, through quantitative time-series model fitting, we have elucidated critical and complementary roles of MinE’s membrane-binding and dimerization domains.

### Model selection and redundant parameters

The AABSM has more parameters than other recently proposed models (e.g. [4, 37]), and so an improved fit over these is not surprising. In fact, as demonstrated by the removed-reaction study, some parameters are clearly redundant. We included them nonetheless to allow for an analysis that highlights which reactions are most important, and used the AIC score to account for this complexity advantage. We note that the SAM has even more parameters than the AABSM but did not fit the data as well, as measured by the *χ*^2^ value. This suggests to us that the AABSM is not just an improvement over earlier models by virtue of additional parameters but because some of those parameters represent reactions that better reflect the underlying biochemistry.

### Variable experimental conditions introduce modelling uncertainty

The oscillation-data and the MinD-dissociation-data experiments were carried out with different membrane compositions and buffer flow rates. We expected parameter values to vary somewhat between the two data sets so we estimated parameters for each experiment independently. As shown in Fig. S9, when we instead fit the two data sets simultaneously, constraining the parameters used for each experiment to be within roughly an order of magnitude of each other, we find that the AABSM can still fit both sets of time-course data nearly as well. The parameter values from this approach are shown in Table S7.

In the removed-reaction study, we found that the removal of some parameters did not impact the quality of the fit significantly after re-optimization of the remaining parameters. In all such cases, when the AIC score dropped for one of the data sets, it rose in the other, indicating that the AABSM fitting has some degrees of freedom that might be eliminated when both data sets are fit simultaneously with the same parameter values for the two experiments. Instead of imposing that parameters be the same in both experiments, which would likely be incorrect given the variability in the experimental conditions, we point out that Fig. S9 shows fits for a continuum of progressively looser constraints. Ultimately, simplifying the AABSM is beyond the scope of what we can do with the time-course data we have and will require the incorporation of more experimental evidence and additional theoretical considerations.

Variability in optimal parameter values both across fits of the AABSM model and its removed-reaction simplifications and across the data sets serves as a warning about the biological accuracy of our parameter values. This combined with the degrees of freedom in the AABSM fitting made it difficult to meaningfully test some of the hypotheses described above through simulation, including the roles of membrane binding and dimerization.

## Methods

### Data

To fit the models, we use *in vitro* data, kindly provided by Ivanov and Mizuuchi [28] (oscillation data) and Vecchiarelli *et al.* [49] (MinD dissociation data). The geometry, space scale, and more deterministic behavior of the *in vitro* assays allows improved opportunities for analysis in comparison with *in vivo* data. Due to these advantages in combination with the constant buffer flow in the experiments and the spatial near-homogeneity of the data, the data fitting problem was reduced from one of fitting each of the PDE models to one of fitting the corresponding ODEs, each with three fewer states (eliminating dynamic variables for concentrations of bulk MinD monomers, bulk MinD dimers, and bulk MinE dimers). In other words, we focused on fitting the reaction kinetics without the complications of spatial dynamics and variable bulk states.

Some preprocessing of Ivanov and Mizuuchi’s data was required. Fluorescence intensities of fluorescently labeled MinD and MinE were not spatially aligned, so we used cross-correlation to align the images, taking advantage of of similarly shaped structures in the two signals. To remove background noise from the aligned data, we used images from before any protein reached the imaging region of the flow cell to calculate an average noise intensity over space and time and subtracted that from all subsequent images. Then we flattened the shifted aligned data to correct for illumination vignetting by multiplying by the reciprocal of an optimally chosen Guassian that we normalized to have a maximum value of 1. The conversion factor from MinE fluorescence intensity to MinE density was not experimentally determined. Using flattened MinD and MinE fluorescent intensities before a significant amount of either protein bound to the supported lipid bilayer (with a slight correction for a small but increasing amount of bound MinD), known buffer concentrations of MinD and MinE, and the depth of the evanescent waves in TIRF microscopy, we calculated conversions from flattened MinD and MinE fluorescent intensities to MinD and MinE molecule densities. We found our conversion for MinD to be in fairly good agreement with Ivanov and Mizuuchi’s estimate.

To extract a time series from Ivanov and Mizuuchi’s data that best represents reaction kinetics only, we looked for a region in the images that was as close to spatially homogeneous as possible, avoiding, for example, regions and time intervals with traveling wave fronts moving through them. To do so, we searched over all disks of 1000 pixels in the preprocessed data for the disk with the least spatial variation in MinD and MinE densities during a single period of the oscillation. To reduce pixel-to-pixel noise in measuring this spatial variation, we smoothed the preprocessed data using values from local, radially-symmetric 2-D linear regressions over disks of 100 pixels. Ultimately, for the model fitting, we used mean values of non-smoothed MinD and MinE densities at each time within the chosen disk as the time series for model fitting. The resulting oscillation data is shown in the top panel of Fig. S1 and in normalized form in the top panels of Figures 2 and 3 (dotted). SEMs were omitted because they were indistinguishable by eye from the means.

Vecchiarelli *et al.*’s data consisted of MinD and MinE time series, both normalized by the concentration of MinD at the beginning of dissociation, and raw MinD time-series data. To calibrate the time-series data, we first aligned normalized MinD data and raw MinD data under best-fitting affine transformations. Once aligned, scalings in each of the best-fitting affine transformations provided us with conversions from normalized units to the arbitrary units (AU) used in the experiments. We calibrated normalized MinD and MinE time-series data for model fitting using the conversion from AU to dimers/*μm*^2^ reported by Vecchiarelli *et al.*. The resulting MinD dissociation data is shown in the bottom panel of Fig. S1 and in normalized form in the bottom panels of Figures 2 and 3 (dotted). We note that, unlike the oscillation data which we calibrated directly from data earlier in the same experiment, Vecchiarelli *et al.* calibrated MinD dissociation data using data from a different experiment of their own with a similar setup, so MinD dissociation data may contain more calibration error than the oscillation data.

### Models

The reactions included in each of the models are displayed in Fig. 6. Below, we describe the modifications made to the Bonny Model [4] as well as the relationships between each of the tested models.

For consistency with buffer-flow experimental conditions, we fixed the bulk concentrations of MinD and MinE (see the Supporting Information Methods from [28] for justification). The original Bonny Model has no spontaneous MinD dissociation which is critical for fitting Vecchiarelli’s MinD dissociation data with MinE absent, so we included reaction **d → D** in the Asymmetric Activation Model. The form of the term we used for this reaction (not mass action) is described and justified in the Supplemental Material.

Because the SAM and AABSM are more complicated models than the AAM, aiming to explain the more recent experimental results that the AAM cannot account for, we also included the CAAM in our model selection process. The CAAM essentially includes the intersection of all reaction terms found in the SAM and AABSM, giving it more scope to account for some of those more recent observations (see the Supplemental Material for more discussion). This additional scope does not allow it to account for the dual role of MinE which is why we tested the SAM and AABSM.

We included a wide range of possible reactions in the SAM and the AABSM. We added all backward reactions except those where MinE dimers spontaneously dissociate from complexes with MinD dimers on the membrane and return to the bulk (we added these to the CAAM as well). Those excluded backward reactions were omitted because residence times of MinE dimers are at least 1.3 times as long as residence times of MinD dimers in all portions of Min-protein traveling waves on a supported lipid bilayer *in vitro* [31]. We also excluded reactions where MinD dimers in **de** complexes return to the bulk in the SAM and where MinD dimers in **ded** complexes return to the bulk in the AABSM, to enforce stability in membrane-stable MinD states. As shown in Table S1, including MinD-membrane dissociation reactions for membrane-stable MinD states in the SAM and AABSM does not change our model selection results.

### Parameter fitting

To find optimal data-fitting numerical solutions for our models, we fit them to the oscillation data and the MinD dissociation data using the homotopy-minimization method for parameter estimation in differential equations [6]. Apart from the constrained optimizations shown in Table S7, we fit models to the two sets of time-course data independently. Details of data fitting are described in the Supplemental Material.

The oscillation data, dissociation data, and matlab scripts that simulate the AABSM in both contexts with optimal parameters are available at https://github.com/ecytryn/Min-biochemistry.

## Acknowledgements

The authors acknowledge funding support from NSERC through a grant to ENC. This research was enabled in part by support provided by Westgrid (https://www.westgrid.ca/), Calcul Quebec (https://www.calculquebec.ca/en/) and Compute Canada (www.computecanada.ca). We thank Kiyoshi Mizuuchi and Anthony Vecchiarelli for sharing fluorescence images and other data from their experiments.

## Supplemental material

### S1 ODE Models

#### S1.1 Model Notation

- *c*_*d*_ is the concentration of membrane-bound MinD dimers.
- *c*_*de*_ is the concentration of MinD-MinE hetero-tetramer complexes on the membrane.
- *c*_*ede*_ is the concentration of MinE-MinD-MinE hetro-hexamer complexes on the membrane.
- *c*_*ded*_ is the concentration of MinD-MinE-MinD hetero-hexamer complexes on the membrane.
- *c*_*e*_ is the concentration of membrane-bound MinE dimers.
- *c*_*D*_ is the constant concentration of bulk MinD dimers.
- *c*_*E*_ is the constant concentration of bulk MinE dimers.
- *ω*_*u,v→x,y*_ denotes the reaction rate of *c*_*u*_ and *c*_*v*_ converting into *c*_*x*_ and *c*_*y*_ for *u, v, x, y* ∈ {*Ø, D, E, d, de, ede, ede, e*}.
- 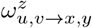 denotes the reaction rate of *c*_*u*_ and *c*_*v*_ converting into *c*_*x*_ and *c*_*y*_ with facilitation by *c*_*z*_ for *u, v, x, y, z* ∈ {*Ø, D, E, d, de, ede, ede, e*}.
- *c*_max_ is the saturation concentration of MinD dimers on the membrane.
- *c*_*s*_ is the half-max concentration constant in the Hill equation modeling the rate of spontaneous MinD-membrane dissociation.
- *n*_*s*_ is the Hill coefficient in the Hill equation modeling the rate of spontaneous MinD-membrane dissociation.
- 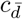 is the constant concentration of MinD dimers that are persistently bound to the membrane. This accounts for the effect of MinD dimers that remain permanently bound to the membrane, as observed in experiments [S28], on spontaneous MinD-membrane dissociation.
- *C*_*d*_ is the constant concentration of MinD monomers that are not accounted for in ODE models, the planar-density of constant bulk MinD monomers illuminated within the depth of the evanescence wave in TIRF microscopy and the concentration of MinD monomers that are persistently bound to the membrane.
- *C*_*e*_ is the constant concentration of MinE monomers that are not accounted for in ODE models, the planar-density of constant bulk MinE monomers illuminated within the depth of the evanescence wave in TIRF microscopy and the concentration of MinE monomers that are persistently bound to the membrane.

#### S1.2 Modeling Spontaneous MinD Dissociation from the Membane

Without the inclusion of spontaneous MinD dissociation from the membrane in models, fitting MinD dissociation data with MinE absent would be meaningless. As the residence time of MinD on the supported lipid bilayer increases from 11 s to at least 40.71 s as the concentration of MinD increases from 0.275 *μ*M to 1.1 *μ*M in the absence of MinE [S31] and the mechanism underlying this concentration-dependent stabilization is not known, we phenomenologically modeled the rate of spontaneous MinD-membrane dissociation by a reverse hill function,

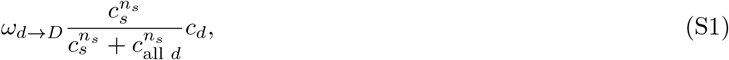

where *ω*_*d→D*_ is the maximum value of the off rate, *c*_*s*_ is the MinD dimer concentration on the membrane at the half-max value of the off rate, *c*_all *d*_ is the concentration of all MinD dimers (in all complexes) on the membrane (see model-specific forms in the differential-equation models below), *c*_*d*_ is the concentration of (MinE-free) membrane-bound MinD dimers, and *n*_*s*_ is the Hill coefficient. Apart from this reaction, **d → D**, all reactions in all models follow the law of mass action.

#### S1.3 Asymmetric Activation Model (AAM)

We define the AAM such that

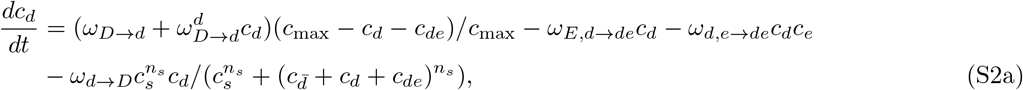

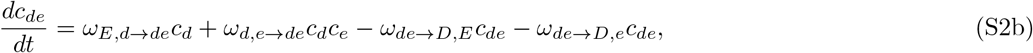

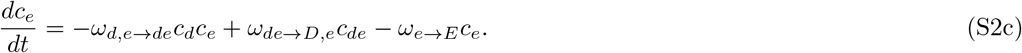

Its MinD monomer density = 2(*c*_*d*_ + *c*_*de*_) + *C*_*d*_, and its MinE monomer density = 2(*c*_*de*_ + *c*_*e*_) + *C*_*e*_. For fitting to the MinD dissociation data with MinE in the flowed buffer, we set 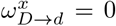 for *x* ∈ {*Ø, d*}. For fitting to the MinD dissociation data with MinE absent, we set *c*_*de*_ = 0, *c*_*e*_ = 0, and all nontrivial rate parameters except *ω*_*d→D*_ equal to 0. We note that *ω*_*D→d*_ and 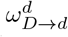 have a multiplicative factor of *c*_*D*_ built into them and *ω*_*E,d→de*_ has a multiplicative factor of *c*_*E*_ built into it.

#### S1.4 Comprehensive Asymmetric Activation Model (CAAM)

The AAM assumes that membrane-bound MinD dimers, but not membrane-bound MinD-MinE tetramers, recruit bulk MinD dimers to bind to the membrane. In MinD and MinE bursts on a supported lipid bilayer *in vitro*, however, increasing the buffer concentration of MinE increases both the maximal net membrane-attachment rate of MinD and the peak membrane density of MinD [S49]. Because a presumed increase of MinE binding to MinD on the membrane does not seem to suppress MinD’s ability to recruit bulk MinD to bind to the membrane, we allow membrane-bound MinD-MinE tetramers to recruit bulk MinD dimers to bind to the membrane 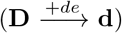 in the CAAM. We know of no model that incorporates membrane-bound MinE facilitated recruitment of bulk MinE to bind to MinE-free MinD on the membrane. Without such, an increasing net membrane-attachment rate of MinE from roughly 5 s until 20 s in the MinD dissociation data would not likely be possible as the density of MinE-free MinD on the membrane, bulk MinE’s substrate, presumably decreases during that time period. Thus, we allow membrane-bound MinD-MinE tetramers and membrane-bound MinE dimers to recruit bulk MinE dimers to bind to membrane-bound MinD dimers 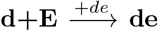 and 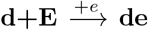 in the CAAM. Additionally, we allow the possibility of some backward reactions in the CAAM, as discussed in the Methods. Including the aforementioned reactions into the AAM, we define the CAAM such that

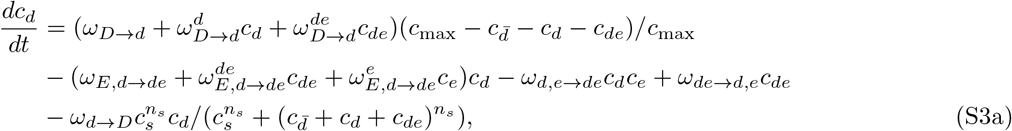

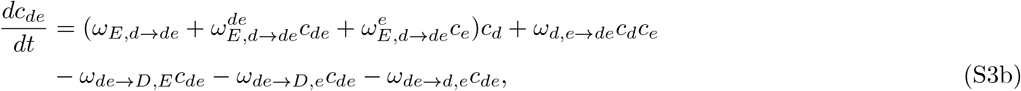

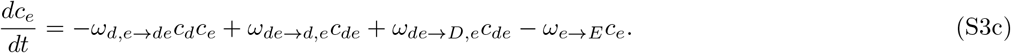

Like the AAM, its MinD monomer density = 2(*c*_*d*_ + *c*_*de*_) + *C*_*d*_, and its MinE monomer density = 2(*c*_*de*_ + *c*_*e*_) + *C*_*e*_. For fitting to the MinD dissociation data with MinE in the flowed buffer, we set 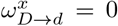 for *x* ∈ {*Ø, d, de*}. For fitting to the MinD dissociation data with MinE absent, like with the AAM, we set *c*_*de*_ = 0, *c*_*e*_ = 0, and all nontrivial rate parameters except *ω*_*d→D*_ equal to 0. We note that 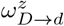 has a multiplicative factor of *c*_*D*_ built into it for *z* ∈ {*Ø, d, de*} and 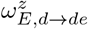 has a multiplicative factor of *c*_*E*_ built into it for *z* ∈ {*Ø, d, de*}.

#### S1.5 Symmetric Activation Model (SAM)

We define the SAM such that

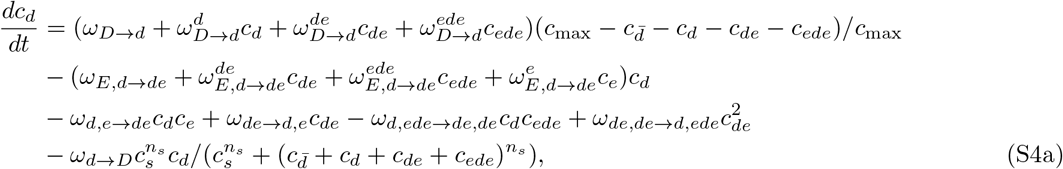

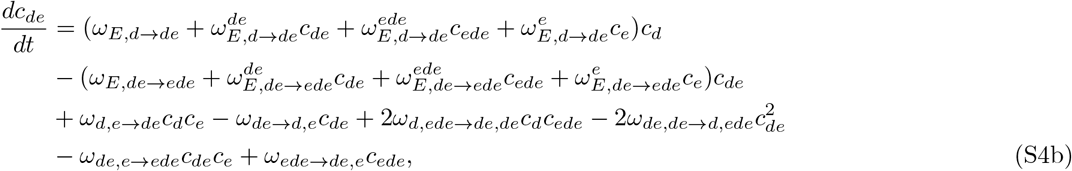

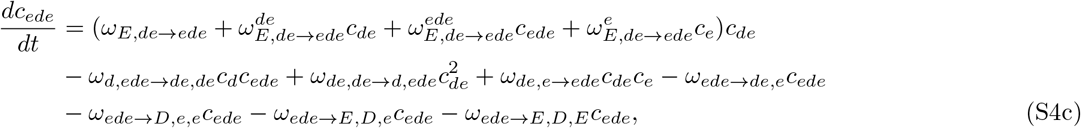

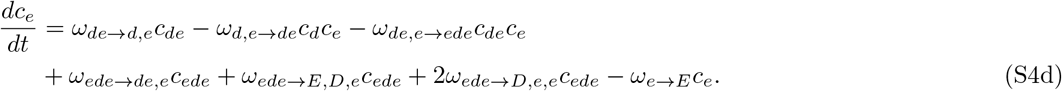

Its MinD monomer density = 2(*c*_*d*_ + *c*_*de*_ + *c*_*ede*_) + *C*_*d*_, and its MinE monomer density = 2(*c*_*de*_ + 2*c*_*ede*_ + *c*_*e*_) + *C*_*e*_. For fitting to the MinD dissociation data with MinE in the flowed buffer, we set 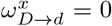 for *x* ∈ {*Ø, d, de, ede*}. For fitting to the MinD dissociation data with MinE absent, we set *c*_*de*_, *c*_*ede*_ = 0, *c*_*e*_ = 0, and all nontrivial rate parameters except *ω*_*d→D*_ equal to 0. We note that 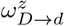 has a multiplicative factor of *c*_*D*_ built into it for *z* ∈ {*Ø, d, de, ede*} and 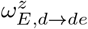 and 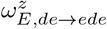 have a multiplicative factor of *c*_*E*_ built into them for *z* ∈ {*Ø, de, ede, e*}.

#### S1.6 Asymmetric Activation with Bridged Stability Model (AABSM)

We define the AABSM such that

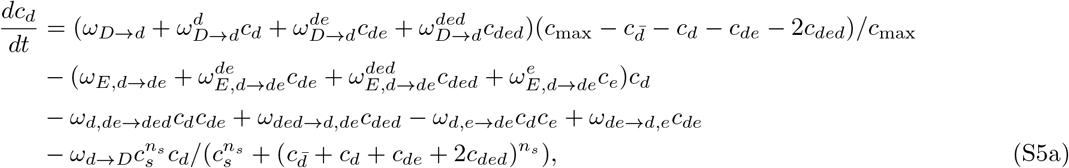

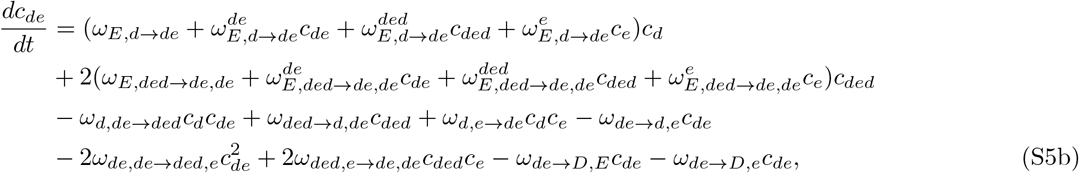

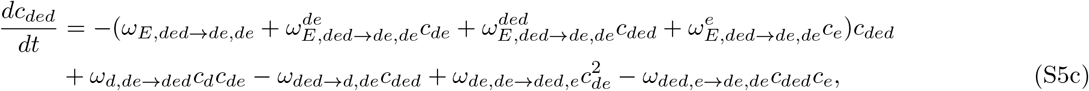

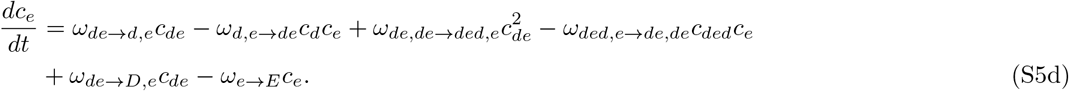

Its MinD monomer density = 2(*c*_*d*_ + *c*_*de*_ + 2*c*_*ded*_) + *C*_*d*_, and its MinE monomer density = 2(*c*_*de*_ + *c*_*ded*_ + *c*_*e*_) + *C*_*e*_. For fitting to the MinD dissociation data with MinE in the flowed buffer, we set 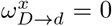 for *x* ∈ {*Ø, d, de, ded*}. For fitting to the MinD dissociation data with MinE absent, we set *c*_*de*_ = 0, *c*_*ded*_ = 0, *c*_*e*_ = 0, and all nontrivial rate parameters except *ω*_*d→D*_ equal to 0. We note that 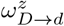 has a multiplicative factor of *c*_*D*_ built into it for *z* ∈ {*Ø, d, de, ded*} and 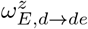 and 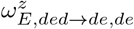 have a multiplicative factor of *c*_*E*_ built into them for *z* ∈ {*Ø, de, ded, e*}.

#### S1.7 The FitzHugh Nagumo Model (FHNM)

This FHNM is simplified model of neuron firing, not the Min system, that we use to test how well an arbitrary model can describe the time-course data. The parameters in the model are *a, b, c, d, I, v*_*l*_, and *v*_*u*_. The FHNM model is defined such that

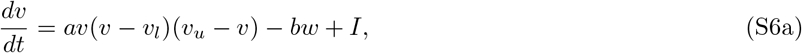

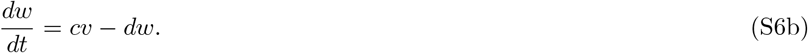

We define its MinD monomer density as *v* + *w* + *C*_*d*_ and its MinE monomer density as *w* + *C*_*e*_. For fitting to the MinD dissociation data with MinE in the flowed buffer, we include all states and all parameters in the model. For fitting to the MinD dissociation data with MinE absent, we set *w* = 0.

### S2 Implementation of the Homotopy-Minimization Method for Data Fitting

We employ the Homotopy-Minimization Method for Parameter Estimation in Differential Equations [S6] to fit our models to the time-course data. In this section, we first define our statistical model for use in the method, then we describe details of method implementation for the AAM, the CAAM, the SAM, and the AABSM. We omit details of the FHNM for brevity, but note that they are similar in nature to those discussed.

#### S2.1 Statistical Model

For the *j*^th^ (of *n*_*y*_) data value *d*_*jk*_ measured at time *t*_*k*_ for *k* ∈ {1, 2 ..., *n*_*t*_(*j*)}, corresponding observable model value *y*_*jk*_, and error *ε*_*jk*_,

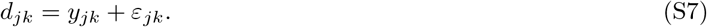

As examples, for just the oscillation data and the AAM, *n*_*y*_ = 2, *d*_1*k*_ is the measured density of MinD monomers at time *t*_*k*_, *d*_2*k*_ is the density of MinE monomers at time *t*_*k*_, *y*_1*k*_ = 2(*c*_*d*_ + *c*_*de*_)+*C*_*d*_ at time *t*_*k*_, and *y*_2*k*_ = 2(*c*_*de*_ + *c*_*e*_)+*C*_*e*_ at time *t*_*k*_. Errors, *ε*_*jk*_, consist of modeling errors and data errors. We find that standard errors of the mean (SEMs) in the oscillation data range between roughly 5 *μm*^−2^ and 30 *μm*^−2^ while the data ranges on the scale of 5·10^3^ *μm*^−2^. We did not extract MinD dissociation data from raw data, so we do not know its SEMs, but we assume that they are also small compared to data ranges. As such, we expect modeling errors to be significantly larger than data errors, and thus expect errors, *ε*_*jk*_, to consist primarily of modeling errors. Modeling errors are inherently not independent nor identically distributed, but without a better a priori distribution, we assume that *ε*_*jk*_ are independent and identically distributed from a normal distribution with a mean of 0, for *j* = 1,..., *n*_*y*_ and *k* = 1, 2 ..., *n*_*t*_(*j*). Ranges of protein densities vary in our time-course data. Thus, to remove bias in fitting from differences in scale, we assume that *ε*_*jk*_ are proportional to 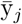, the range of *d*_*jk*_ for *k* = 1, 2 ..., *n*_*t*_(*j*), for *j* = 1,..., *n*_*y*_ and *k* = 1, 2 ..., *n*_*t*_(*j*). Therefore, collectively, we assume that

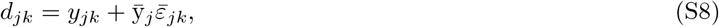

where 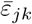 are independent and identically distributed from a normal distribution with a mean of 0 and a variance of 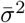, 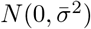, for *j* = 1,..., *n*_*y*_ and *k* = 1, 2 ..., *n*_*t*_(*j*). Thus, the likelihood of *y*_*jk*_ given *d*_*jk*_ and 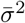 for *j* = 1,..., *n*_*y*_ and *k* = 1, 2 ..., *n*_*t*_(*j*) is given by

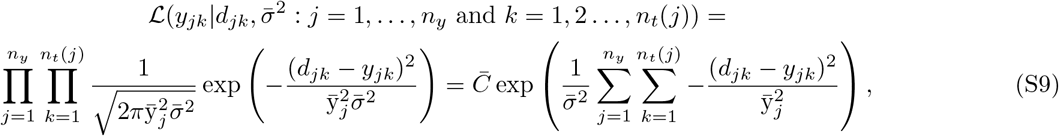

for constant 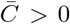. The values of *y*_*jk*_ for *j* = 1,..., *n*_*y*_ and *k* = 1, 2 ..., *n*_*t*_(*j*) that maximize the likelihood are those that minimize

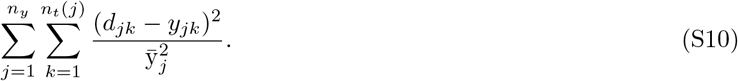

Thus, we measure the difference in observable model values from time-course data by the sum of weighted squared residuals in Equation S10.

#### S2.2 Defining Functionals, *r*_*y*_(p, x) and *r*_Δ*x*_(p, x), for the Homotopy-Minimization Method

As described above, we measure the difference in observable model values from time-course data by the sum of weighted squared residuals in Equation S10. Thus, we define *r*_*y*_(**p**, **x**), the measure of data fitting as defined in [S6], such that

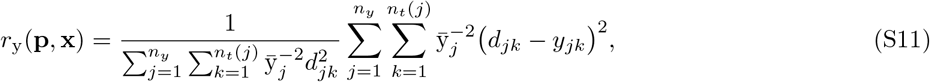

where we normalize by 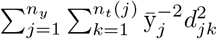 to match the scale of *r*_*y*_(**p**, **x**) in Equation 2.6 of [S6]. We do not penalize deviations from interpolated data in our fitting, so we define 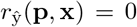, for 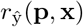 as described in Section 2.6 of [S6].

For consistency with the notation in [S6], we define *x*_1_ = *c*_*d*_, *x*_2_ = *c*_*de*_, and *x*_3_ = *c*_*e*_ (*n*_*x*_ = 3) for the AAM and the CAAM; *x*_1_ = *c*_*d*_, *x*_2_ = *c*_*de*_, *x*_3_ = *c*_*ede*_, and *x*_4_ = *c*_*e*_ (*n*_*x*_ = 4) for the SAM; and *x*_1_ = *c*_*d*_, *x*_2_ = *c*_*de*_, *x*_3_ = *c*_*ded*_, and *x*_4_ = *c*_*e*_ (*n*_*x*_ = 4) for the AABSM. We define *r*_Δ*x*_(**p**, **x**), the measure of satisfying a numerical solution to a differential-equation model as defined in [S6], as in Equation 2.13a of [S6] and discretize models using a Simpson’s method finite difference, a finite difference with fourth order accuracy. Thus, in r_Δ*x*_(**p**, **x**),

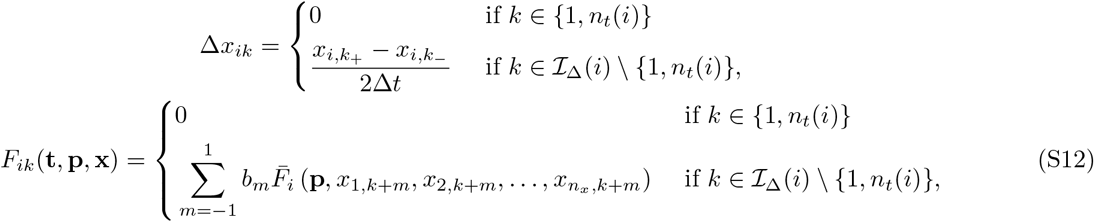

where *k*_+_ is the index above *k* in 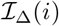 (the index set of the numerical discretization), *k*_−_ is the index below *k* in 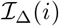, Δt is the grid spacing in {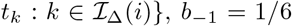, b_0_ = 4/6, b_1_ = 1/6, and 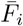 is the right-hand side of the ODE for *x*_*i*_, for *i* = 1,..., *n*_*x*_. For smoothing penalties in *r*_Δ*x*_(**p**, **x**), *s*_*i*_(**x**) as defined in [S6], we set *α*_*i*_ = 1, *β*_*i*_ = 10^2^, and *γ*_*i*_ = 2, for *i* = 1,..., *n*_*x*_, as implemented in [S6] to insignificantly modify r_Δ*x*_(**p**, **x**) with a smooth set of state values and to strongly penalize r_Δ*x*_(**p**, **x**) with a jagged set of state values.

#### S2.3 Domain Restrictions on States and Parameters

We restrict parameters and state values in models to be consistent with biological assumptions and certain experimental measurements. Rate parameters, 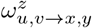 for *u, v, x, y, z* ∈ {*Ø, D, E, d, de, ede, ded, e*}, are only biologically relevant if nonnegative, and we assume that reactions are not overly fast, with rate parameters exceeding 10 units. Thus, we restrict rate parameters such that

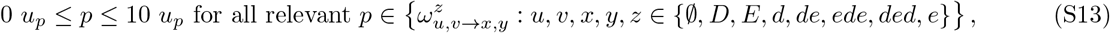

where *u*_*p*_ is the units of parameter *p*. Additionally, for the SAM and the AABSM including MinD-membrane-dissociation reactions of membrane-stable MinD states, as discussed in Table S1, to impose each model’s core assumption, we restrict the rate of MinD dimer-membrane dissociation to be no faster in the membrane-stable MinD state than the membrane-unstable MinD state:

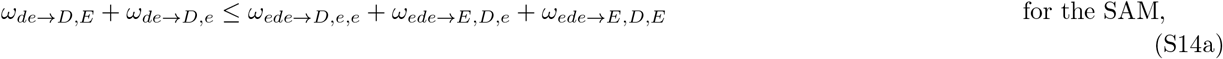

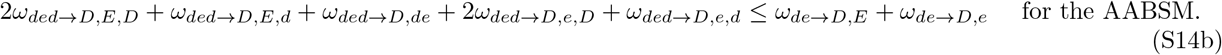

The parameter *c*_max_, which is only pertinent for fitting the oscillation data, dictates the maximum concentration of membrane-bound MinD dimers and is necessarily no less than half the range of the MinD monomer density in the time course, 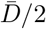. Additionally, we assume that *c*_max_ is on the scale of 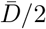, so we bound *c*_max_ above by 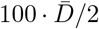. Thus, we restrict *c*_max_ such that

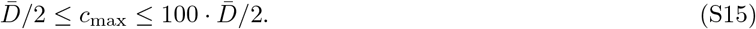

We also assume that *c*_*s*_, the half-max concentration constant in the Hill equation modeling the rate of spontaneous MinD-membrane dissociation, is on the scale of 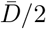, so we bound *c*_*s*_ above by 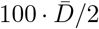. The Hill equation modeling the rate of spontaneous MinD-membrane dissociation can be undefined if *c*_*s*_ = 0, so we bound *c*_*s*_ below by 1 *μm*^−2^. Thus, we restrict *c*_*s*_ such that

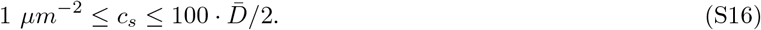

The Hill coefficient in the Hill equation modeling the rate of spontaneous MinD-membrane dissociation, *n*_*s*_, is necessarily nonnegative, and we assume that it does not exceed a (biologically very large) value of 10, so we restrict *n*_*s*_ such that

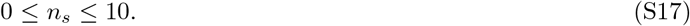

The constant planar-density of bulk MinD monomers and the constant concentration of persistent membrane-bound MinD monomers, *C*_*d*_, is necessarily nonnegative and no greater than the minimum MinD monomer density in each MinD time course, *D*_min_, so we restrict *C*_*d*_ such that

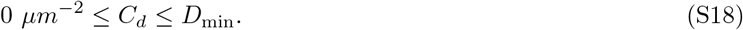

Similarly, for the oscillation data, *C*_*e*_, the constant planar-density of bulk MinE monomers and the constant concentration of persistent membrane-bound MinE monomers, is necessarily nonnegative and no greater than the minimum MinE monomer density in the time course, *E*_min_. The density of MinE is 0 *μm*^−2^ at the start of the MinD dissociation data with MinE in the flowed buffer, so *E*_min_ is not a meaningful upper bound of *C*_*e*_ for the time course. As such, we restrict *C*_*e*_ such that

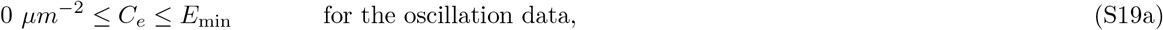

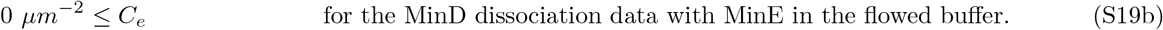

The constant concentration of persistent membrane-bound MinD dimers, 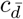, is necessarily nonnegative and no greater than half the value of *C*_*d*_, which includes the concentration of persistent membrane-bound MinD monomers in it, so we restrict 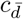 such that

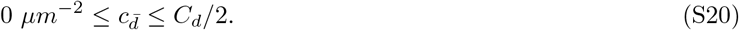

Concentrations *c*_*d*_, *c*_*de*_, *c*_*ede*_, *c*_*ded*_, and *c*_*e*_ are only biologically relevant if nonnegative. Thus, we restrict *c*_*d*_, *c*_*de*_, *c*_*ede*_, *c*_*ded*_, and *c*_*e*_ to nonnegative values:

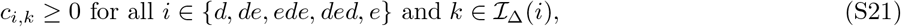

where *c*_*d,k*_, *c*_*de,k*_, *c*_*ede,k*_, *c*_*ded,k*_, and *c*_*e,k*_ are the values of *c*_*d*_, *c*_*de*_, *c*_*ede*_, *c*_*ded*_, and *c*_*e*_ at the *k*^th^ index of the numerical discretization.

All aforementioned restrictions on parameters and state vales can be written as a collection of linear inequalities. Thus, during accelerated descent, as implemented in overlapping-niche descent of the Homotopy-Minimization Method, we employ projection using Dykstra’s method as discussed in Section C.2.3 of [S6]. In doing so, we choose a small relative termination tolerance, *ε*_*c*_ = 10^−6^, and a smaller absolute termination tolerance, 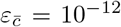. To avoid overly slowing accelerated descent from a large number of projections, we prematurely terminate accelerated descent if the number of iterations in Dykstra’s method exceeds 10^4^.

#### S2.4 Generating Random Parameter and State Values

Following the procedure of overlapping-niche descent in the Homotopy-Minimization Method, as discussed in Section C.1 of [S6], we randomly generate parameters and state values initially and in random offspring. Given no prior parameter-value estimates, we randomly generate rate parameters over a broad range of scales in accordance with their bounds, Equation S13:

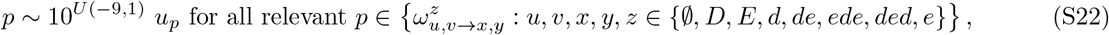

where *U(a, b)* is the uniform probability distribution over the interval (*a, b*) and *u*_*p*_ is the units of parameter *p*.

We expect *c*_max_, which is only pertinent for fitting the oscillation data, to be within one or two orders of magnitude of half the range of the MinD monomer density in the time course, 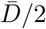. Thus, in accordance with the bounds on *c*_max_, Equation S15, we randomly generate *c*_max_ such that

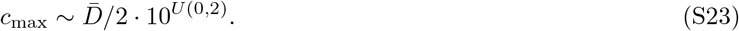

Similarly, we expect *c*_*s*_ to be within one or two orders of magnitude of 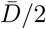. Thus, in accordance with the bounds on *c*_*s*_, Equation S16, we randomly generate *c*_*s*_ such that

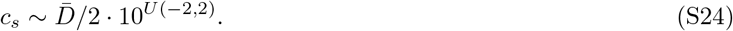

We randomly generate *n*_*s*_, *C*_*d*_, *C*_*e*_, and 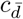 uniformly over their respective bounds, Equations S17, S18, S19, and S20:

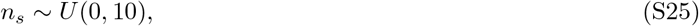

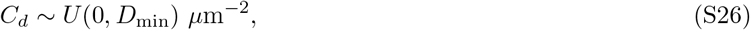

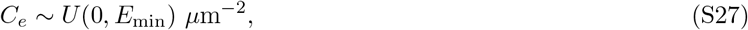

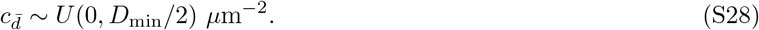

We choose random state values to match time-course data exactly. Thus, for the oscillation data and the MinD dissociation data with MinE in the flowed buffer, MinD monomer density *D*, MinE monomer density *E*, *D*_2_ = (*D* − *C*_*d*_)/2, and *E*_2_ = (*E* − *C*_*e*_)/2, we generate random state values for the AAM and the CAAM with *c*_*e*_ as a free state such that

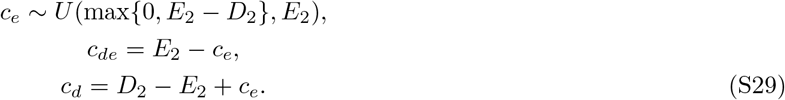

Similarly, we generate random state values for the SAM with *c*_*ede*_ and *c*_*e*_ as free states such that

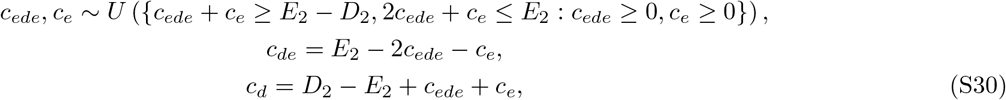

and we generate random state values for the AABSM with *c*_*ded*_ and *c*_*e*_ as free states such that

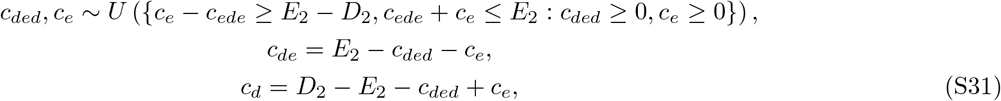

where *U*({·}) is the uniform probability distribution over the set {·}. Additionally, for the MinD dissociation data with MinE absent, we generate random state values for the AAM, the CAAM, the SAM, and the AABSM with no free states such that

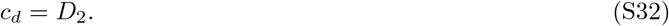

#### S2.5 Random Perturbation and Selection

During the generation of sexual offspring in overlapping-niche descent of the Homotopy-Minimization Method, as described in Section C.1 of [S6], for each sexual offspring, we define the standard deviation of perturbation, 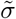, such that

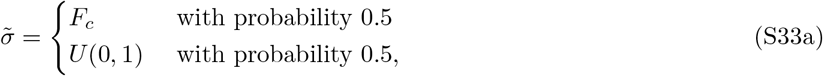

where *F*_*c*_ is a measure of convergence defined below and *U(a, b)* is the uniform probability distribution over the interval (*a, b*). Then, for parameter *p* in a sexual offspring inherited from individual (**p**_*g,i,j*_, **x**_*g,i,j*_), we perturb the value of the inherited parameter, 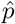, such that

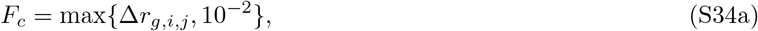

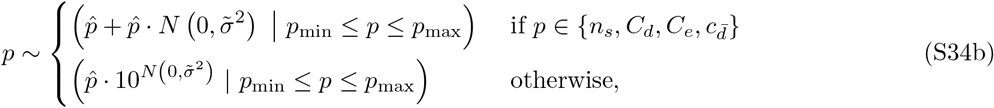

where Δ*r*_*g,i,j*_ is the measure of convergence in the *j*^th^ parent space of the *i*^th^ niche in generation *g* as defined in Equation C.1 of [S6], *N*(*μ, σ*^2^) is the normal distribution with mean *μ* and variance *σ*^2^, and *p*_min_ and *p*_max_ are the restricted lower and upper bounds of parameter *p* as discussed in Section S2.3. In the standard deviation of perturbation, 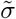, max{Δ*r*_*g,i,j*_, 10^−2^} ensures diminishing but significant perturbations in 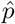 as the algorithm converges, and *U*(0, 1) allows for a wide range of perturbations in 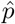. Similarly, for state value *x* in a sexual offspring inherited from individual (**p**_*g,i,j*_, **x**_*g,i,j*_), we perturb the value of the inherited state value, 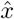, such that

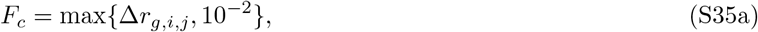

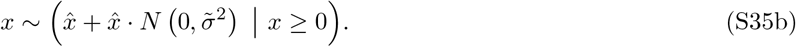

During selection in overlapping-niche descent, as described in Section C.1 of [S6], we choose the natural default value for the selection strength, *q*_fit_ = 1.

#### S2.6 Remaining Details for Implementation of the Homotopy-Minimization Method

We choose values of *λ* to define niches in overlapping-niche descent of the Homotopy-Minimization Method as in Section 3.4.3 of [S6]. We choose parents and offspring in overlapping-niche descent as in Section E.1.2 of [S6]. For initial gradient scaling values, *s*_*i,*0_ for *i* in the indexed set of all parameters and state values, as described in Section C.2.1 of [S6], we choose s_*i,*0_ = 10^−12^ for all *i* corresponding to parameters to ensure that projections onto restrictions involving multiple parameters are initially defined, and we choose *s*_*i,*0_ = 0 for all *i* corresponding to state values. We choose *n*_max_, *ε*_*σ*_, *ε*_*r*_, *n*_max_, 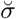, *m*_pro_, 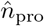, 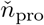, and *ε*_Δ*r*_ in overlapping-niche descent as in Section E.1.5 of [S6]. Finally, we calculate confidence intervals for parameters using bootstrapping, as in Section 3.4.4 of [S6].

We computed genetic-algorithm components of the Homotopy-Minimization Method using *MATLAB*, and we computed accelerated-descent components of the Homotopy-Minimization Method in parallel using *C++* on the Calcul Québec server Guillimin, the Compute Canada servers Cedar and Graham, and the WestGrid server Orcinus.

#### S2.7 Numerical-Discretization Refinements

Initially, we fit each model to the time-course data using the data grid as the numerical-discretization grid. For some of the models, the numerical solution that fit the time-course data best would contain spurious oscillations. In such cases, we refined the numerical-discretization grid, increasing *ρ*, the ratio of the number of (evenly spaced) grid points in the numerical discretization to the number of data points in the time course, by whole numbers until the spurious oscillations disappeared. As such, for the AAM, *ρ* = 3 for the oscillation data, and *ρ* = 2 for the MinD dissociation data; for the CAAM, *ρ* = 1 for the oscillation data, and *ρ* = 3 for the MinD dissociation data; for the SAM, *ρ* = 1 for both sets of the time-course data; for the AABSM, *ρ* = 1 for both sets of the time-course data; and for the FHNM, *ρ* = 3 for both sets of the time-course data.

### S3 Data and Model Fits Not Shown in the Main Text

**Figure S1:**
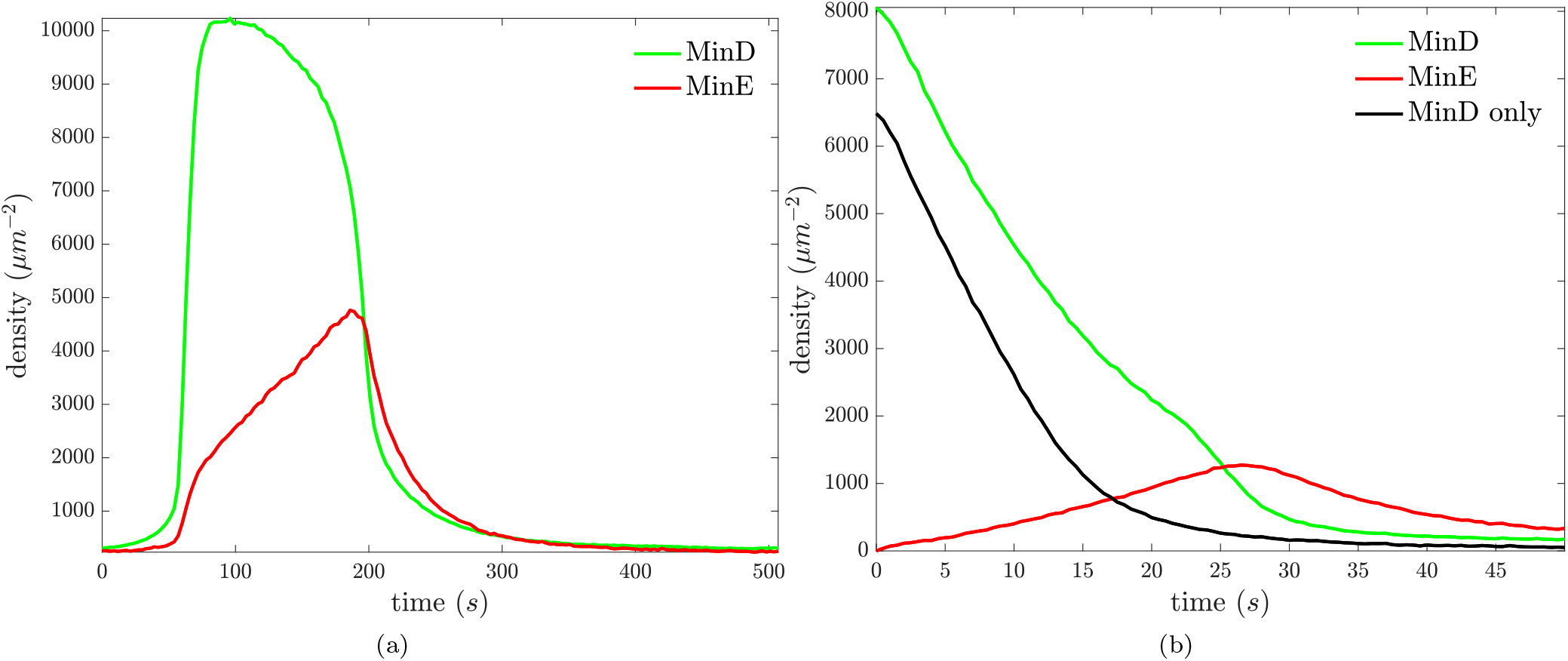
The oscillation data is shown in (a), and MinD dissociation data is shown in (b).

**Figure S2:**
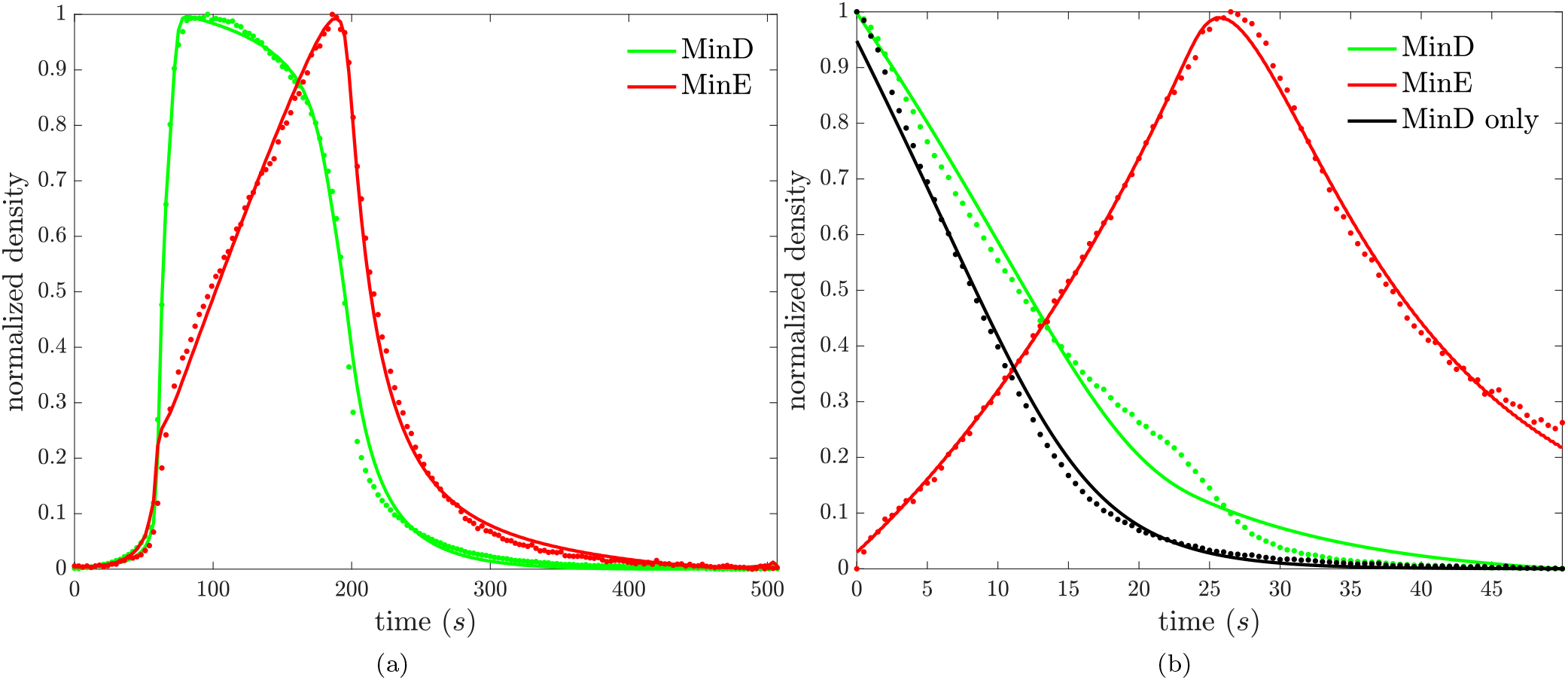
For the CAAM, the fit to the oscillation data is shown in (a), and the fit to the MinD dissociation data is shown in (b). Fits are solid, and the data are dotted.

**Figure S3:**
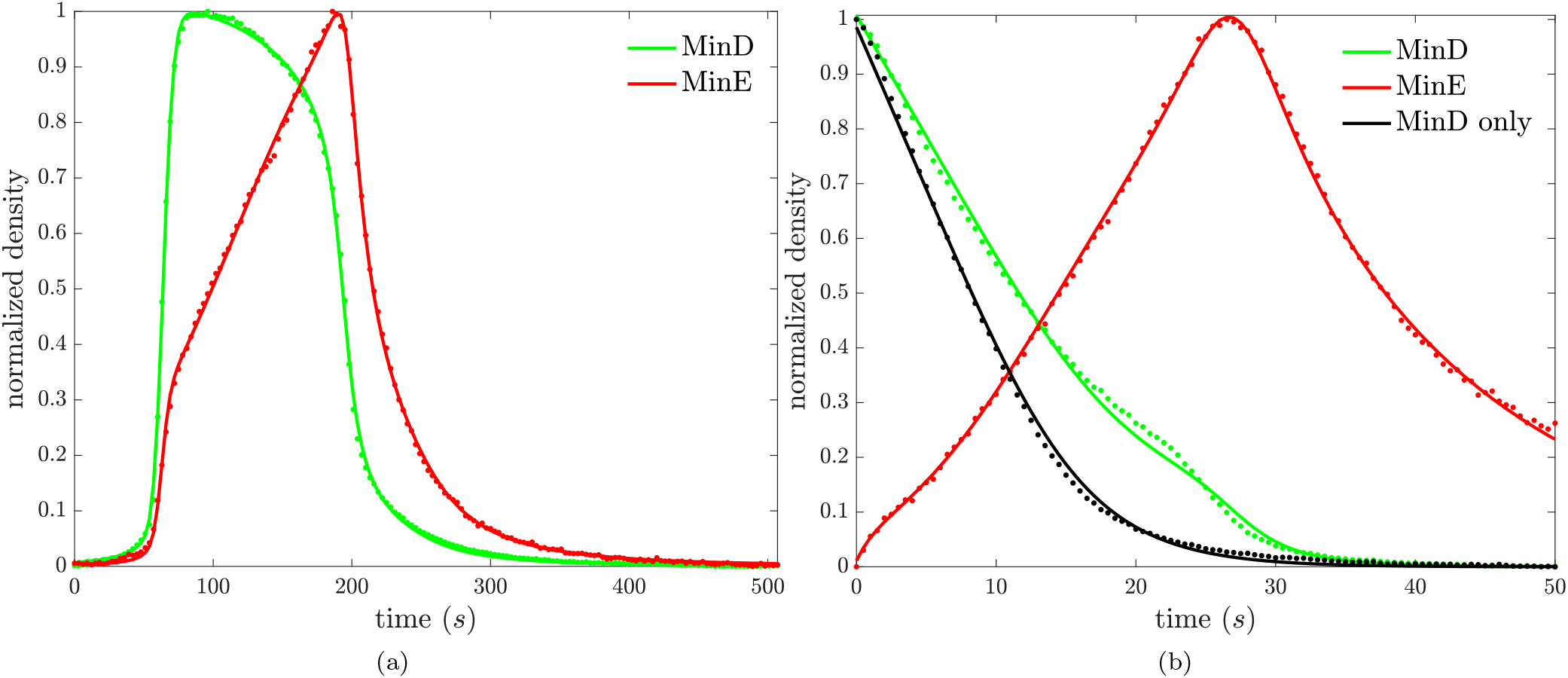
For the SAM, the fit to the oscillation data is shown in (a), and the fit to the MinD dissociation data is shown in (b). Fits are solid, and the data are dotted.

**Figure S4:**
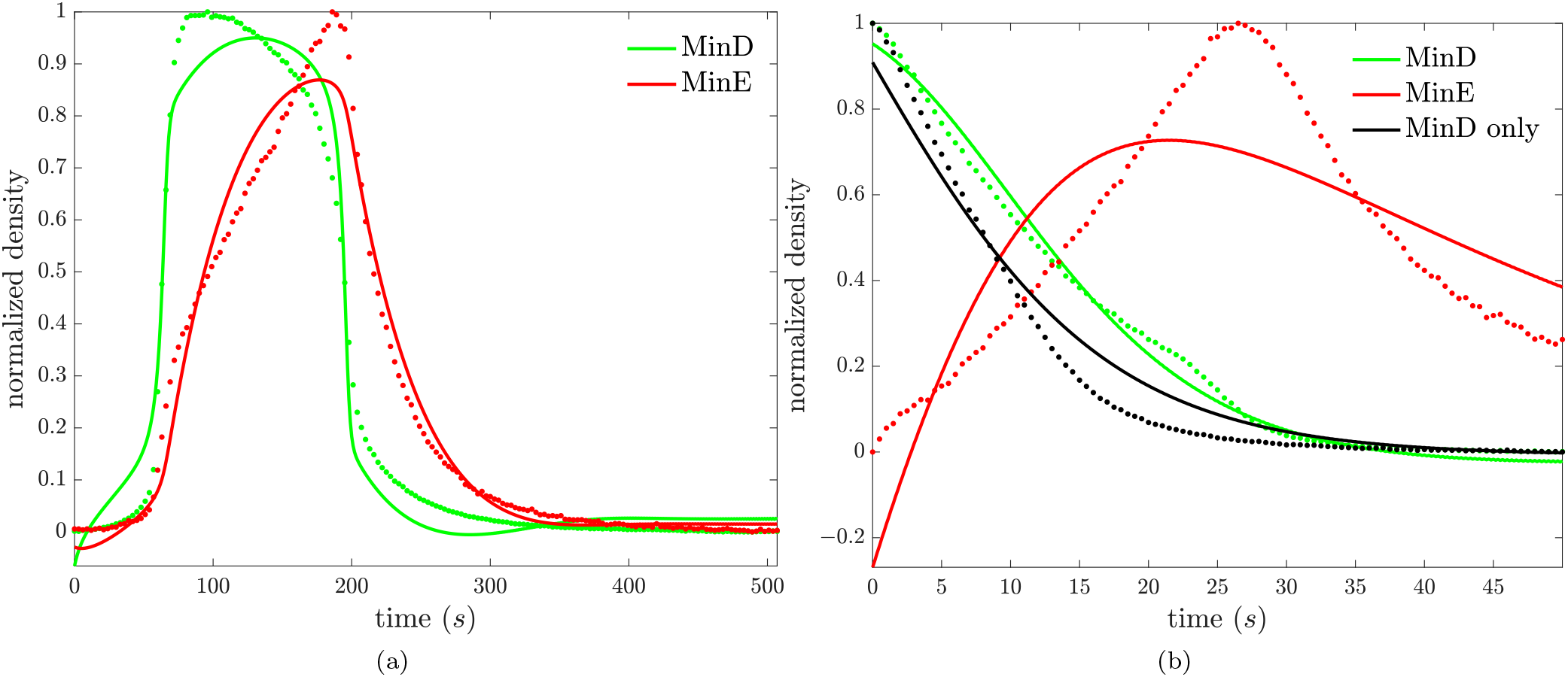
For the FHNM, the fit to the oscillation data is shown in (a), and the fit to the MinD dissociation data is shown in (b). Fits are solid, and the data are dotted.

### S4 State Values from Model Fits Not Shown in the Main Text

**Figure S5:**
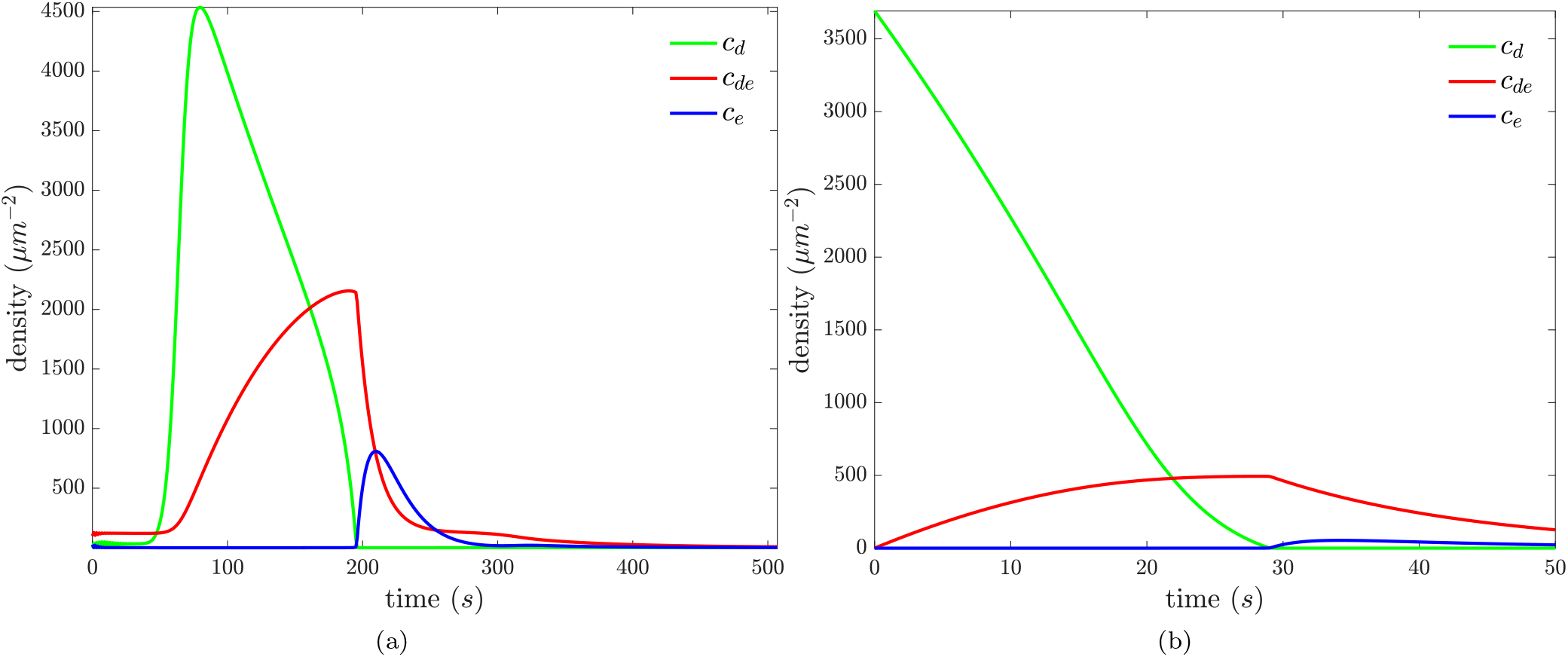
For the AAM, state values from the fit to the oscillation data are shown in (a), and state values from the fit to the MinD dissociation data with MinE in the flowed buffer are shown in (b).

**Figure S6:**
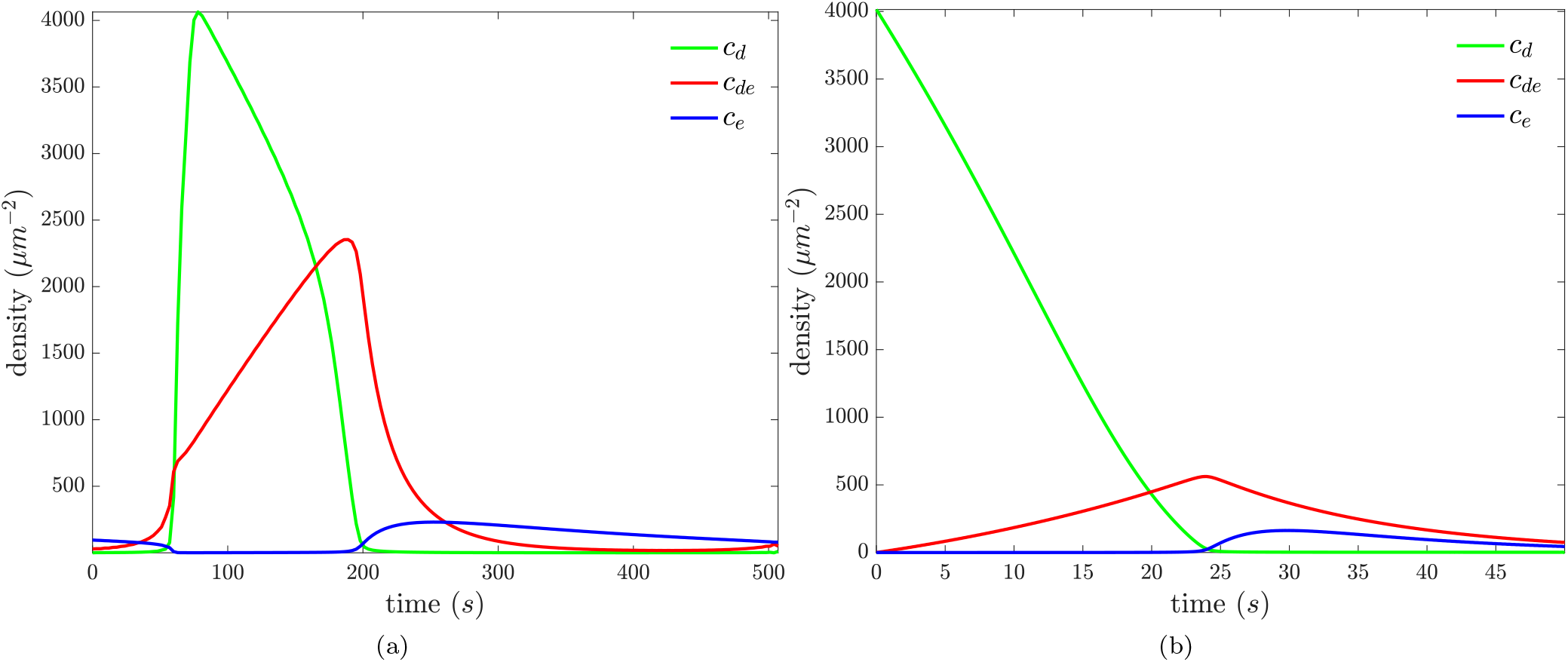
For the CAAM, state values from the fit to the oscillation data are shown in (a), and state values from the fit to the MinD dissociation data with MinE in the flowed buffer are shown in (b).

**Figure S7:**
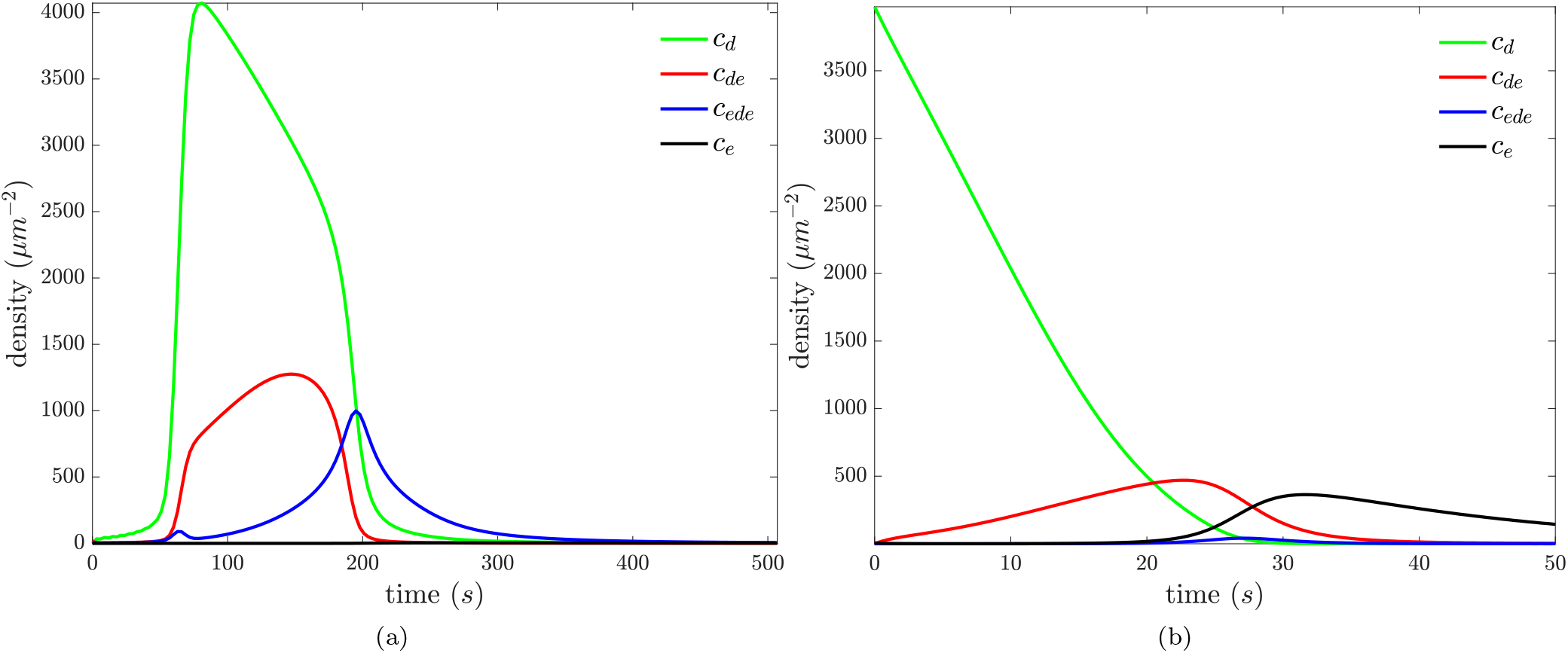
For the SAM, state values from the fit to the oscillation data are shown in (a), and state values from the fit to the MinD dissociation data with MinE in the flowed buffer are shown in (b).

**Figure S8:**
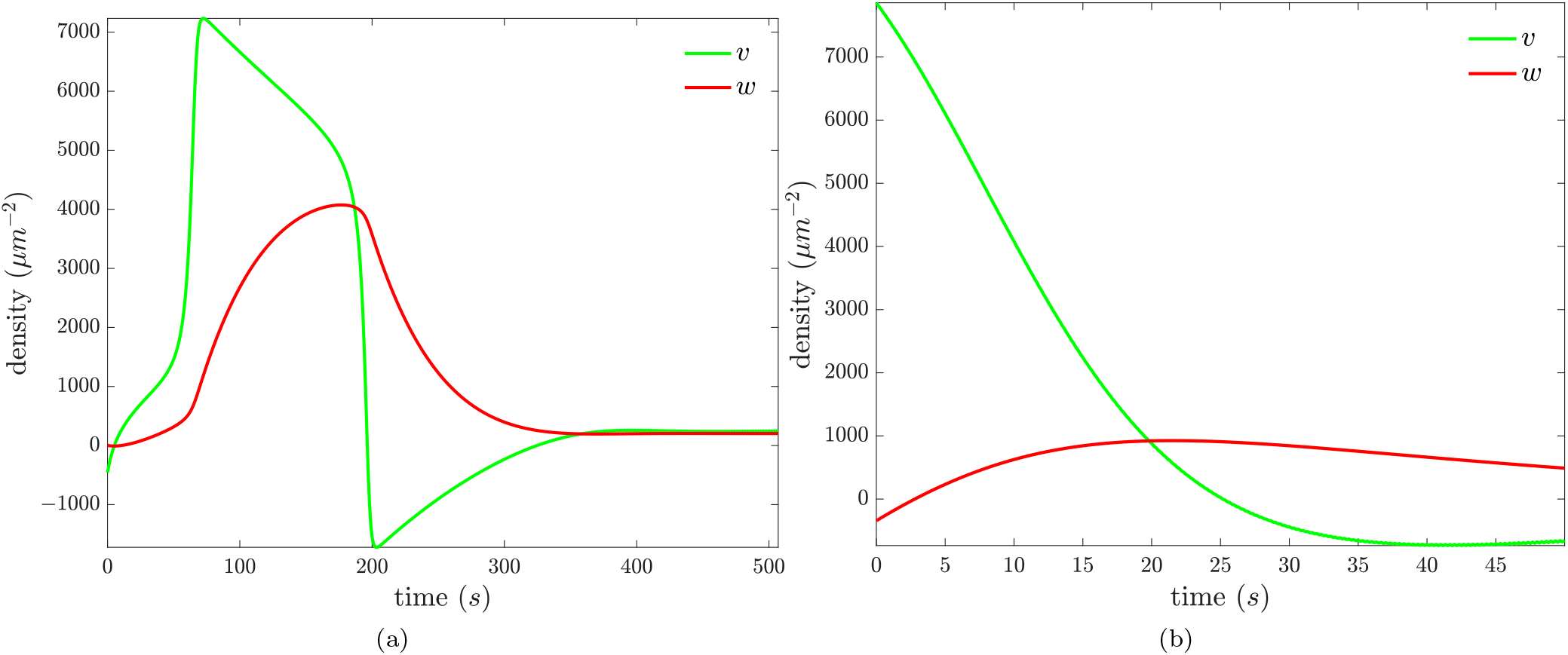
For the FHNM, state values from the fit to the oscillation data are shown in (a), and state values from the fit to the MinD dissociation data with MinE in the flowed buffer are shown in (b).

### S5 The Simultaneous Fitting of Both Data Sets with Parameter Constraints and the Inclusion of MinD-Membrane-Dissociation Reactions of Membrane-Stable MinD States in Models

**Figure S9:**
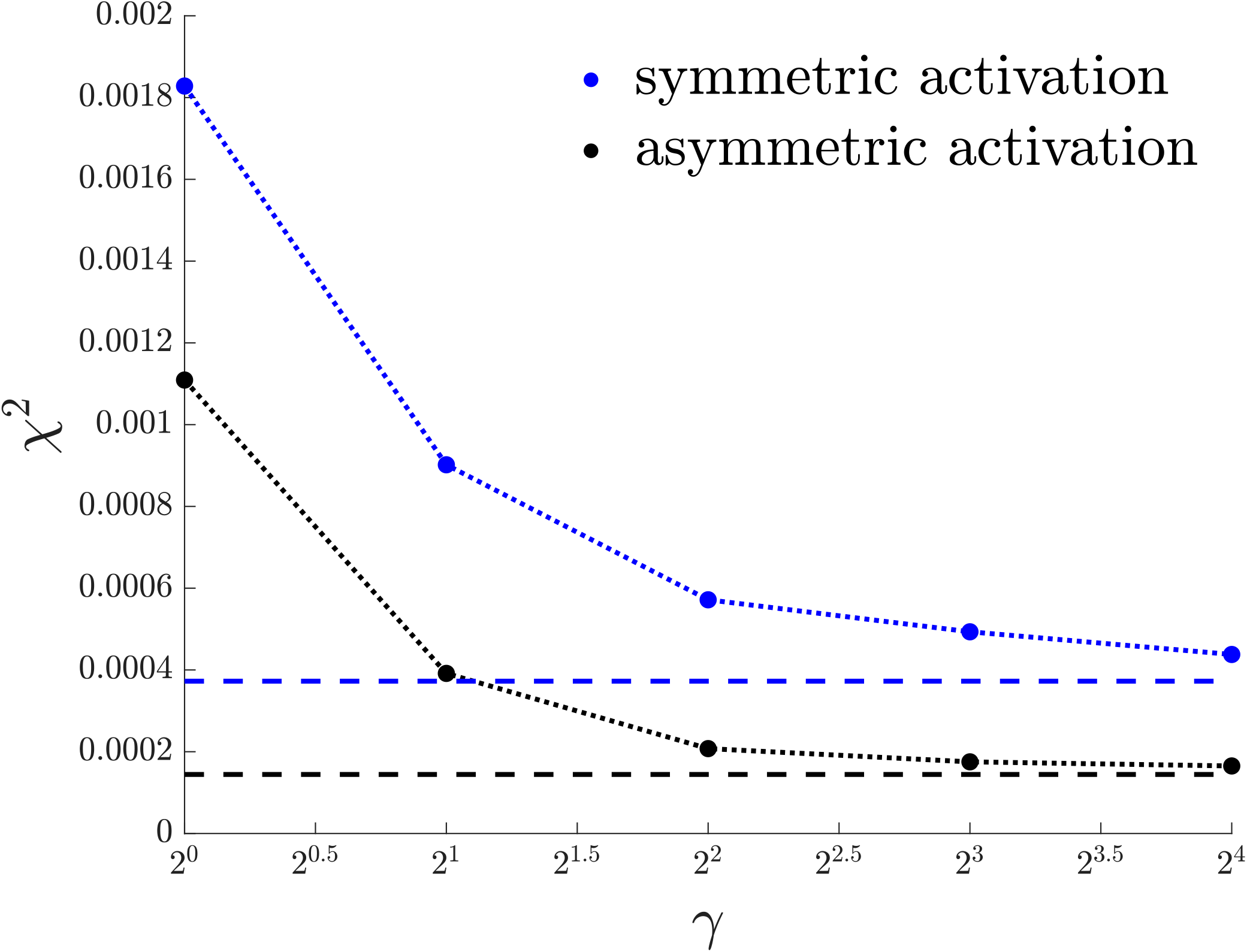
The simultaneous fitting of both data sets with parameter constraints. For each parameter *p* that is nontrivial in both the model of the oscillation data and the model of the MinD dissociation data with MinE in the flowed buffer, except for constant-concentration parameters *C*_*d*_, *C*_*e*_, and 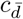, we constrain *p* in the fit of the oscillation data to be less than or equal to *γp* in the fit of the MinD dissociation data, and we constrain *p* in the fit of the MinD dissociation data to be less than or equal to *γp* in the fit of the oscillation data, for *γ* = 1, 2, 4, 8, 16. The resulting *χ*^2^ values from the fits are shown as dots. The *χ*^2^ values from unconstrained optimizations, in which *γ* = *∞*, are shown as flat dashed lines. When parameters across the data sets are constrained to be within roughly an order of magnitude of each other, the models, most notably the AABSM, can still recapitulate both data sets almost as well as without constraints. In fitting, we take into account differences in buffer MinE concentrations – 1.36*μM* for the oscillation data and 2.5*μM* for the MinD dissociation data with MinE in the flowed buffer – by multiplying rate parameters of bulk MinE binding reactions, 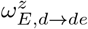 and 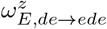 for *z* ∈ {*Ø, de, ede, e*} in the SAM and 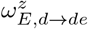 and 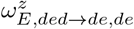 for *z* ∈ {*Ø, de, ded, e*} in the AABSM, which otherwise have a multiplicative factor of *c*_*E*_ built into them, by 1.36*μM* or 2.5*μM*. This removes the multiplicative factor of *c*_*E*_ from the rate parameters of bulk MinE binding reactions and instead incorporates a conversion factor from bulk MinE monomers to dimers in them, under the assumption that all bulk MinE is stable in the dimer state. Parameter estimates for the AABSM with *γ* = 16 are shown in Table S7.

**Table S1:** Model Comparison with and without MinD-membrane-dissociation reactions of membrane-stable MinD states. An asterisk (*) indicates that the model includes MinD-membrane-dissociation reactions of the membrane-stable MinD state, the reactions de → D+E and de → D+e for the SAM and the reactions ded → D+E+D, ded → D+E+d, ded → D+de, ded → D+e+D, and ded → D+e+d for the AABSM. *χ*^2^ is the weighted sum of squared residuals, and AIC is the Akaike information criterion. 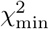 is the minimum *χ*^2^ value amongst the models, and AIC_min_ is the minimum AIC value amongst the models. The AABSM without the inclusion of MinD-membrane-dissociation reactions of the membrane-stable MinD state fits the oscillation data and the MinD dissociation data better than the SAM, even with the inclusion of MinD-membrane-dissociation reactions of the membrane-stable MinD state in the SAM. Including MinD-membrane-dissociation reactions of the membrane-stable MinD state in the AABSM does not appreciably improve its fit to the oscillation data and does not improve its fit to the MinD dissociation data. We include this table as justification for the omission of the membrane-stable MinD dissociation reactions from the SAM and AABSM as mentioned in the Methods.

|  | Oscillation Data |  | MinD Dissociation Data |  |
| --- | --- | --- | --- | --- |
| | $\chi^2/\chi_{\min}^2$ | AIC - AIC<sub>min</sub> | $\chi^2/\chi_{\min}^2$ | AIC - AIC<sub>min</sub> |
| SAM | 1.9 | $2.1 \cdot 10^2$ | 3.2 | $3.6 \cdot 10^2$ |
| SAM* | 1.5 | $1.3 \cdot 10^2$ | 3.2 | $3.5 \cdot 10^2$ |
| AABSM | 1.1 | $1.2 \cdot 10^1$ | 1.0 | 0 |
| AABSM* | 1 | 0 | 1 | 9.3 |

### S6 Parameter Estimates from Fits

We provide parameter estimates from our fits of the models to the time-course data below. In them, if a parameter is blank, it was not included in the fitting. All parameters with non-blank and non-zero values were included in the parameter counts of AIC scores. Because the time-course data is not very noisy, fitting errors consist primarily of modeling errors, and modeling errors rather than independent and identically distributed Gaussian errors were redistributed during the calculation of confidence intervals by bootstrapping. As such, the confidence intervals shown are not very biologically informative and are often smaller for models that fit the time-course data better, but we provide them for reference anyway.

**Table S2:** Parameters from the fits of the AAM to the oscillation data and the MinD dissociation data.

|  | Oscillation Data |  | MinD Dissociation Data |  |  |
| --- | --- | --- | --- | --- | --- |
| Parameter | Value | 95% Confidence Interval | Value | 95% Confidence Interval | Units |
| $C_d$ | $3.0 \cdot 10^2$ | $[2.8 \cdot 10^2, 3.0 \cdot 10^2]$ | 0 | $[0, 0]$ | $\mu m^{-2}$ |
| $C_d$ (w/o MinE) | | | $5.1 \cdot 10^1$ | $[0, 5.1 \cdot 10^1]$ | $\mu m^{-2}$ |
| $C_e$ | $2.3 \cdot 10^2$ | $[1.6 \cdot 10^2, 2.3 \cdot 10^2]$ | 0 | $[0, 3.9 \cdot 10^1]$ | $\mu m^{-2}$ |
| $c_{\bar{d}}$ | 0 | $[0, 2.4 \cdot 10^1]$ | 0 | $[0, 0]$ | $\mu m^{-2}$ |
| $c_{\bar{d}}$ (w/o MinE) | | | 0 | $[0, 2.5 \cdot 10^1]$ | $\mu m^{-2}$ |
| $c_{\max}$ | $5.5 \cdot 10^3$ | $[5.4 \cdot 10^3, 5.6 \cdot 10^3]$ | | | $\mu m^{-2}$ |
| $c_s$ | $9.4 \cdot 10^1$ | $[5.8 \cdot 10^1, 1.6 \cdot 10^2]$ | $1.5 \cdot 10^3$ | $[1.3 \cdot 10^3, 1.9 \cdot 10^3]$ | $\mu m^{-2}$ |
| $n_s$ | 7.6 | $[2.9, 1.0 \cdot 10^1]$ | 2.3 | $[1.9, 3.2]$ | |
| $\omega_{D \rightarrow d}$ | $1.0 \cdot 10^1$ | $[8.5, 1.0 \cdot 10^1]$ | | | $\mu m^{-2} s^{-1}$ |
| $\omega_{D \rightarrow d}^d$ | $2.4 \cdot 10^{-1}$ | $[2.2 \cdot 10^{-1}, 2.5 \cdot 10^{-1}]$ | | | $s^{-1}$ |
| $\omega_{E, d \rightarrow de}$ | $6.1 \cdot 10^{-3}$ | $[5.7 \cdot 10^{-3}, 6.5 \cdot 10^{-3}]$ | $1.0 \cdot 10^{-2}$ | $[1.0 \cdot 10^{-2}, 1.1 \cdot 10^{-2}]$ | $s^{-1}$ |
| $\omega_{d, e \rightarrow de}$ | $3.2 \cdot 10^{-1}$ | $[3.1 \cdot 10^{-1}, 3.3 \cdot 10^{-1}]$ | 9.9 | $[9.7, 1.0 \cdot 10^1]$ | $\mu m^2 s^{-1}$ |
| $\omega_{d \rightarrow D}$ | $1.0 \cdot 10^1$ | $[1.5, 1.0 \cdot 10^1]$ | $2.1 \cdot 10^{-1}$ | $[1.7 \cdot 10^{-1}, 2.5 \cdot 10^{-1}]$ | $s^{-1}$ |
| $\omega_{de \rightarrow D, E}$ | $1.7 \cdot 10^{-3}$ | $[1.0 \cdot 10^{-3}, 2.4 \cdot 10^{-3}]$ | 0 | $[0, 3.1 \cdot 10^{-3}]$ | $s^{-1}$ |
| $\omega_{de \rightarrow D, e}$ | $7.5 \cdot 10^{-2}$ | $[6.9 \cdot 10^{-2}, 7.7 \cdot 10^{-2}]$ | $6.5 \cdot 10^{-2}$ | $[6.0 \cdot 10^{-2}, 7.0 \cdot 10^{-2}]$ | $s^{-1}$ |
| $\omega_{e \rightarrow E}$ | $6.2 \cdot 10^{-2}$ | $[5.5 \cdot 10^{-2}, 7.0 \cdot 10^{-2}]$ | $4.3 \cdot 10^{-1}$ | $[3.8 \cdot 10^{-1}, 4.5 \cdot 10^{-1}]$ | $s^{-1}$ |

**Table S3:** Parameters from the fits of the CAAM to the oscillation data and the MinD dissociation data.

|  | Oscillation Data |  | MinD Dissociation Data |  |  |
| --- | --- | --- | --- | --- | --- |
| Parameter | Value | 95% Confidence Interval | Value | 95% Confidence Interval | Units |
| $C_d$ | $2.6 \cdot 10^2$ | $[2.3 \cdot 10^2, 3.0 \cdot 10^2]$ | $1.2 \cdot 10^1$ | $[0, 2.7 \cdot 10^1]$ | $\mu m^{-2}$ |
| $C_d$ (w/o MinE) | | | $5.1 \cdot 10^1$ | $[1.7 \cdot 10^{-1}, 5.1 \cdot 10^1]$ | $\mu m^{-2}$ |
| $C_e$ | 0 | $[0, 2.0 \cdot 10^1]$ | $3.8 \cdot 10^1$ | $[1.4 \cdot 10^1, 2.4 \cdot 10^1]$ | $\mu m^{-2}$ |
| $c_{\bar{d}}$ | $5.3 \cdot 10^1$ | $[5.7, 1.1 \cdot 10^2]$ | 6.0 | $[0, 1.2 \cdot 10^1]$ | $\mu m^{-2}$ |
| $c_{\bar{d}}$ (w/o MinE) | | | 0 | $[0, 1.7 \cdot 10^1]$ | $\mu m^{-2}$ |
| $c_{\max}$ | $6.7 \cdot 10^3$ | $[6.7 \cdot 10^3, 6.7 \cdot 10^3]$ | | | $\mu m^{-2}$ |
| $c_s$ | $4.5 \cdot 10^3$ | $[4.5 \cdot 10^3, 4.5 \cdot 10^3]$ | $1.7 \cdot 10^3$ | $[1.4 \cdot 10^3, 1.9 \cdot 10^3]$ | $\mu m^{-2}$ |
| $n_s$ | 4.0 | $[4.0, 4.1]$ | 1.7 | $[1.5, 2.1]$ | |
| $\omega_{D \rightarrow d}$ | 0 | $[0, 6.9 \cdot 10^{-2}]$ | | | $\mu m^{-2} s^{-1}$ |
| $\omega_{D \rightarrow d}^d$ | 4.2 | $[4.2, 4.3]$ | | | $s^{-1}$ |
| $\omega_{D \rightarrow d}^{de}$ | $7.0 \cdot 10^{-2}$ | $[6.3 \cdot 10^{-2}, 7.5 \cdot 10^{-2}]$ | | | $s^{-1}$ |
| $\omega_{E, d \rightarrow de}$ | $3.2 \cdot 10^{-3}$ | $[2.5 \cdot 10^{-3}, 3.5 \cdot 10^{-3}]$ | $3.8 \cdot 10^{-3}$ | $[3.1 \cdot 10^{-3}, 4.0 \cdot 10^{-3}]$ | $s^{-1}$ |
| $\omega_{E, d \rightarrow de}^{de}$ | 0 | $[0, 8.5 \cdot 10^{-7}]$ | $3.8 \cdot 10^{-6}$ | $[5.7 \cdot 10^{-7}, 8.7 \cdot 10^{-6}]$ | $\mu m^2 s^{-1}$ |
| $\omega_{E, d \rightarrow de}^e$ | $3.1 \cdot 10^{-2}$ | $[2.9 \cdot 10^{-2}, 3.2 \cdot 10^{-2}]$ | $5.9 \cdot 10^{-2}$ | $[5.8 \cdot 10^{-2}, 6.4 \cdot 10^{-2}]$ | $\mu m^2 s^{-1}$ |
| $\omega_{d, e \rightarrow de}$ | $4.1 \cdot 10^{-3}$ | $[3.8 \cdot 10^{-3}, 4.4 \cdot 10^{-3}]$ | 6.4 | $[6.4, 6.4]$ | $\mu m^2 s^{-1}$ |
| $\omega_{d \rightarrow D}$ | 2.6 | $[2.6, 2.6]$ | $2.0 \cdot 10^{-1}$ | $[1.8 \cdot 10^{-1}, 2.4 \cdot 10^{-1}]$ | $s^{-1}$ |
| $\omega_{de \rightarrow D, E}$ | $9.6 \cdot 10^{-2}$ | $[9.1 \cdot 10^{-2}, 1.0 \cdot 10^{-1}]$ | $2.9 \cdot 10^{-7}$ | $[0, 5.4 \cdot 10^{-3}]$ | $s^{-1}$ |
| $\omega_{de \rightarrow D, e}$ | $1.3 \cdot 10^{-2}$ | $[1.2 \cdot 10^{-2}, 1.4 \cdot 10^{-2}]$ | $7.7 \cdot 10^{-2}$ | $[7.2 \cdot 10^{-2}, 7.7 \cdot 10^{-2}]$ | $s^{-1}$ |
| $\omega_{de \rightarrow d, e}$ | 0 | $[0, 3.3 \cdot 10^{-4}]$ | 7.1 | $[7.1, 7.1]$ | $s^{-1}$ |
| $\omega_{e \rightarrow E}$ | $4.7 \cdot 10^{-3}$ | $[4.3 \cdot 10^{-3}, 5.0 \cdot 10^{-3}]$ | $3.3 \cdot 10^{-1}$ | $[3.2 \cdot 10^{-1}, 3.3 \cdot 10^{-1}]$ | $s^{-1}$ |

**Table S4:** Parameters from the fits of the SAM to the oscillation data and the MinD dissociation data.

|  | Oscillation Data |  | MinD Dissociation Data |  |  |
| --- | --- | --- | --- | --- | --- |
| Parameter | Value | 95% Confidence Interval | Value | 95% Confidence Interval | Units |
| $C_d$ | $3.0 \cdot 10^2$ | $[2.8 \cdot 10^2, 3.0 \cdot 10^2]$ | $1.7 \cdot 10^2$ | $[6.6 \cdot 10^1, 1.3 \cdot 10^2]$ | $\mu m^{-2}$ |
| $C_d$ (w/o MinE) | | | $5.1 \cdot 10^1$ | $[1.2 \cdot 10^1, 5.0 \cdot 10^1]$ | $\mu m^{-2}$ |
| $C_e$ | $2.3 \cdot 10^2$ | $[2.1 \cdot 10^2, 2.3 \cdot 10^2]$ | 1.1 | $[0, 4.6]$ | $\mu m^{-2}$ |
| $c_{\bar{d}}$ | $1.5 \cdot 10^2$ | $[1.3 \cdot 10^2, 1.5 \cdot 10^2]$ | $1.0 \cdot 10^2$ | $[3.2 \cdot 10^1, 6.5 \cdot 10^1]$ | $\mu m^{-2}$ |
| $c_{\bar{d}}$ (w/o MinE) | | | 0 | $[0, 0]$ | $\mu m^{-2}$ |
| $c_{\max}$ | $5.2 \cdot 10^3$ | $[5.2 \cdot 10^3, 5.2 \cdot 10^3]$ | | | $\mu m^{-2}$ |
| $c_s$ | $4.5 \cdot 10^2$ | $[4.2 \cdot 10^2, 4.7 \cdot 10^2]$ | $1.8 \cdot 10^3$ | $[1.5 \cdot 10^3, 2.0 \cdot 10^3]$ | $\mu m^{-2}$ |
| $n_s$ | 3.9 | $[3.2, 4.4]$ | 1.6 | $[1.4, 1.9]$ | |
| $\omega_{D \rightarrow d}$ | 0 | $[0, 1.8 \cdot 10^{-1}]$ | | | $\mu m^{-2} s^{-1}$ |
| $\omega_{D \rightarrow d}^d$ | $2.5 \cdot 10^{-1}$ | $[2.3 \cdot 10^{-1}, 2.6 \cdot 10^{-1}]$ | | | $s^{-1}$ |
| $\omega_{D \rightarrow d}^{de}$ | $7.8 \cdot 10^{-1}$ | $[7.3 \cdot 10^{-1}, 8.5 \cdot 10^{-1}]$ | | | $s^{-1}$ |
| $\omega_{D \rightarrow d}^{ede}$ | $7.9 \cdot 10^{-1}$ | $[7.8 \cdot 10^{-1}, 8.0 \cdot 10^{-1}]$ | | | $s^{-1}$ |
| $\omega_{E,d \rightarrow de}$ | $1.2 \cdot 10^{-3}$ | $[2.2 \cdot 10^{-4}, 1.5 \cdot 10^{-3}]$ | $3.6 \cdot 10^{-5}$ | $[0, 1.4 \cdot 10^{-3}]$ | $s^{-1}$ |
| $\omega_{E,d \rightarrow de}^{de}$ | 0 | $[0, 7.5 \cdot 10^{-7}]$ | $5.3 \cdot 10^{-5}$ | $[3.1 \cdot 10^{-5}, 5.5 \cdot 10^{-5}]$ | $\mu m^2 s^{-1}$ |
| $\omega_{E,d \rightarrow de}^e$ | $9.3 \cdot 10^{-1}$ | $[9.0 \cdot 10^{-1}, 9.5 \cdot 10^{-1}]$ | $1.7 \cdot 10^{-3}$ | $[8.4 \cdot 10^{-4}, 2.4 \cdot 10^{-3}]$ | $\mu m^2 s^{-1}$ |
| $\omega_{E,d \rightarrow de}^{ede}$ | 0 | $[0, 3.2 \cdot 10^{-6}]$ | $1.7 \cdot 10^{-5}$ | $[0, 5.8 \cdot 10^{-3}]$ | $\mu m^2 s^{-1}$ |
| $\omega_{E,de \rightarrow ede}$ | 0 | $[0, 4.7 \cdot 10^{-5}]$ | $7.5 \cdot 10^{-6}$ | $[0, 5.0 \cdot 10^{-4}]$ | $s^{-1}$ |
| $\omega_{E,de \rightarrow ede}^{de}$ | 0 | $[0, 4.4 \cdot 10^{-7}]$ | $9.3 \cdot 10^{-6}$ | $[0, 1.8 \cdot 10^{-5}]$ | $\mu m^2 s^{-1}$ |
| $\omega_{E,de \rightarrow ede}^e$ | 9.9 | $[9.9, 1.0 \cdot 10^1]$ | $1.8 \cdot 10^{-12}$ | $[0, 1.5 \cdot 10^{-4}]$ | $\mu m^2 s^{-1}$ |
| $\omega_{E,de \rightarrow ede}^{ede}$ | $5.3 \cdot 10^{-9}$ | $[0, 7.0 \cdot 10^{-6}]$ | $3.1 \cdot 10^{-3}$ | $[2.3 \cdot 10^{-3}, 3.4 \cdot 10^{-3}]$ | $\mu m^2 s^{-1}$ |
| $\omega_{d,e \rightarrow de}$ | $1.4 \cdot 10^{-4}$ | $[1.1 \cdot 10^{-4}, 1.6 \cdot 10^{-4}]$ | $1.9 \cdot 10^{-4}$ | $[1.1 \cdot 10^{-4}, 4.6 \cdot 10^{-4}]$ | $\mu m^2 s^{-1}$ |
| $\omega_{d,ede \rightarrow de,de}$ | $2.3 \cdot 10^{-4}$ | $[2.3 \cdot 10^{-4}, 2.3 \cdot 10^{-4}]$ | $3.9 \cdot 10^{-4}$ | $[0, 2.0 \cdot 10^{-3}]$ | $\mu m^2 s^{-1}$ |
| $\omega_{d \rightarrow D}$ | $2.8 \cdot 10^{-1}$ | $[2.7 \cdot 10^{-1}, 3.2 \cdot 10^{-1}]$ | $2.1 \cdot 10^{-1}$ | $[1.9 \cdot 10^{-1}, 2.3 \cdot 10^{-1}]$ | $s^{-1}$ |
| $\omega_{de,de \rightarrow d,ede}$ | $3.1 \cdot 10^{-5}$ | $[2.9 \cdot 10^{-5}, 3.2 \cdot 10^{-5}]$ | $6.1 \cdot 10^{-8}$ | $[0, 3.3 \cdot 10^{-6}]$ | $\mu m^2 s^{-1}$ |
| $\omega_{de,e \rightarrow ede}$ | $3.7 \cdot 10^{-3}$ | $[3.5 \cdot 10^{-3}, 4.0 \cdot 10^{-3}]$ | $1.2 \cdot 10^{-3}$ | $[1.1 \cdot 10^{-3}, 1.2 \cdot 10^{-3}]$ | $\mu m^2 s^{-1}$ |
| $\omega_{de \rightarrow d,e}$ | 0 | $[0, 3.2 \cdot 10^{-7}]$ | $4.9 \cdot 10^{-4}$ | $[0, 1.0 \cdot 10^{-2}]$ | $s^{-1}$ |
| $\omega_{e \rightarrow E}$ | $4.1 \cdot 10^{-3}$ | $[4.1 \cdot 10^{-3}, 4.4 \cdot 10^{-3}]$ | $6.2 \cdot 10^{-2}$ | $[6.0 \cdot 10^{-2}, 6.4 \cdot 10^{-2}]$ | $s^{-1}$ |
| $\omega_{ede \rightarrow D,e,e}$ | $8.7 \cdot 10^{-5}$ | $[8.1 \cdot 10^{-5}, 9.4 \cdot 10^{-5}]$ | 1.0 | $[9.2 \cdot 10^{-1}, 1.2]$ | $s^{-1}$ |
| $\omega_{ede \rightarrow E,D,E}$ | $7.9 \cdot 10^{-1}$ | $[7.8 \cdot 10^{-1}, 8.0 \cdot 10^{-1}]$ | $2.5 \cdot 10^{-1}$ | $[2.0 \cdot 10^{-1}, 3.0 \cdot 10^{-1}]$ | $s^{-1}$ |
| $\omega_{ede \rightarrow E,D,e}$ | $2.0 \cdot 10^{-4}$ | $[1.8 \cdot 10^{-4}, 2.1 \cdot 10^{-4}]$ | $7.2 \cdot 10^{-1}$ | $[6.8 \cdot 10^{-1}, 8.0 \cdot 10^{-1}]$ | $s^{-1}$ |
| $\omega_{ede \rightarrow de,e}$ | $1.7 \cdot 10^{-4}$ | $[1.6 \cdot 10^{-4}, 1.8 \cdot 10^{-4}]$ | 1.1 | $[1.0, 1.2]$ | $s^{-1}$ |

**Table S5:** Parameters from the fits of the AABSM to the oscillation data and the MinD dissociation data.

| Parameter | Oscillation Data |  | MinD Dissociation Data |  | Units |
| --- | --- | --- | --- | --- | --- |
|  | Value | 95% Confidence Interval | Value | 95% Confidence Interval |  |
| $C_d$ | $6.5 \cdot 10^1$ | $[5.6 \cdot 10^1, 7.4 \cdot 10^1]$ | $1.7 \cdot 10^2$ | $[6.3 \cdot 10^1, 1.1 \cdot 10^2]$ | $\mu m^{-2}$ |
| $C_d$ (w/o MinE) | | | $5.1 \cdot 10^1$ | $[1.8 \cdot 10^1, 4.5 \cdot 10^1]$ | $\mu m^{-2}$ |
| $C_e$ | 5.4 | [1.4, 9.7] | 0 | [0, 7.3] | $\mu m^{-2}$ |
| $c_{\bar{d}}$ | $1.9 \cdot 10^1$ | $[1.5, 2.6 \cdot 10^1]$ | $8.3 \cdot 10^1$ | $[3.1 \cdot 10^1, 5.4 \cdot 10^1]$ | $\mu m^{-2}$ |
| $c_{\bar{d}}$ (w/o MinE) | | | 0 | [0, 1.1] | $\mu m^{-2}$ |
| $c_{\max}$ | $5.3 \cdot 10^3$ | $[5.2 \cdot 10^3, 5.3 \cdot 10^3]$ | | | $\mu m^{-2}$ |
| $c_s$ | $4.9 \cdot 10^2$ | $[4.6 \cdot 10^2, 5.3 \cdot 10^2]$ | $2.2 \cdot 10^3$ | $[2.0 \cdot 10^3, 2.3 \cdot 10^3]$ | $\mu m^{-2}$ |
| $n_s$ | 9.1 | $[5.4, 1.0 \cdot 10^1]$ | 1.9 | [1.7, 2.1] | |
| $\omega_{D \rightarrow d}$ | 0 | $[0, 7.3 \cdot 10^{-2}]$ | | | $\mu m^{-2} s^{-1}$ |
| $\omega_{D \rightarrow d}^d$ | $3.5 \cdot 10^{-1}$ | $[3.4 \cdot 10^{-1}, 3.5 \cdot 10^{-1}]$ | | | $s^{-1}$ |
| $\omega_{D \rightarrow d}^{de}$ | $2.0 \cdot 10^{-1}$ | $[2.0 \cdot 10^{-1}, 2.0 \cdot 10^{-1}]$ | | | $s^{-1}$ |
| $\omega_{D \rightarrow d}^{ded}$ | $4.0 \cdot 10^{-1}$ | $[3.9 \cdot 10^{-1}, 4.0 \cdot 10^{-1}]$ | | | $s^{-1}$ |
| $\omega_{E,d \rightarrow de}$ | $4.0 \cdot 10^{-3}$ | $[3.5 \cdot 10^{-3}, 4.1 \cdot 10^{-3}]$ | $3.6 \cdot 10^{-3}$ | $[3.1 \cdot 10^{-3}, 4.1 \cdot 10^{-3}]$ | $s^{-1}$ |
| $\omega_{E,d \rightarrow de}^{de}$ | $1.3 \cdot 10^{-10}$ | $[0, 1.2 \cdot 10^{-6}]$ | $7.5 \cdot 10^{-5}$ | $[0, 2.2 \cdot 10^{-4}]$ | $\mu m^2 s^{-1}$ |
| $\omega_{E,d \rightarrow de}^{ded}$ | 0 | $[0, 5.0 \cdot 10^{-7}]$ | $3.6 \cdot 10^{-6}$ | $[0, 1.3 \cdot 10^{-5}]$ | $\mu m^2 s^{-1}$ |
| $\omega_{E,d \rightarrow de}^e$ | 1.0 | [1.0, 1.0] | $4.0 \cdot 10^{-1}$ | $[3.6 \cdot 10^{-1}, 4.4 \cdot 10^{-1}]$ | $\mu m^2 s^{-1}$ |
| $\omega_{E,ded \rightarrow de,de}$ | $2.3 \cdot 10^{-6}$ | $[0, 4.6 \cdot 10^{-4}]$ | $7.7 \cdot 10^{-2}$ | $[6.4 \cdot 10^{-2}, 8.0 \cdot 10^{-2}]$ | $s^{-1}$ |
| $\omega_{E,ded \rightarrow de,de}^{de}$ | $1.4 \cdot 10^{-5}$ | $[1.4 \cdot 10^{-5}, 1.5 \cdot 10^{-5}]$ | $6.6 \cdot 10^{-4}$ | $[5.6 \cdot 10^{-4}, 7.4 \cdot 10^{-4}]$ | $\mu m^2 s^{-1}$ |
| $\omega_{E,ded \rightarrow de,de}^{ded}$ | $1.6 \cdot 10^{-5}$ | $[1.5 \cdot 10^{-5}, 1.6 \cdot 10^{-5}]$ | $6.4 \cdot 10^{-9}$ | $[0, 3.7 \cdot 10^{-5}]$ | $\mu m^2 s^{-1}$ |
| $\omega_{E,ded \rightarrow de,de}^e$ | $8.5 \cdot 10^{-2}$ | $[8.5 \cdot 10^{-2}, 8.6 \cdot 10^{-2}]$ | $4.5 \cdot 10^{-3}$ | $[3.8 \cdot 10^{-3}, 5.3 \cdot 10^{-3}]$ | $\mu m^2 s^{-1}$ |
| $\omega_{d,de \rightarrow ded}$ | $7.2 \cdot 10^{-5}$ | $[7.0 \cdot 10^{-5}, 7.3 \cdot 10^{-5}]$ | $1.1 \cdot 10^{-1}$ | $[1.1 \cdot 10^{-1}, 1.1 \cdot 10^{-1}]$ | $\mu m^2 s^{-1}$ |
| $\omega_{d,e \rightarrow de}$ | $3.0 \cdot 10^{-1}$ | $[3.0 \cdot 10^{-1}, 3.1 \cdot 10^{-1}]$ | 3.4 | [3.4, 3.4] | $\mu m^2 s^{-1}$ |
| $\omega_{d \rightarrow D}$ | $2.5 \cdot 10^{-1}$ | $[2.4 \cdot 10^{-1}, 2.6 \cdot 10^{-1}]$ | $1.9 \cdot 10^{-1}$ | $[1.8 \cdot 10^{-1}, 2.0 \cdot 10^{-1}]$ | $s^{-1}$ |
| $\omega_{de,de \rightarrow ded,e}$ | $1.5 \cdot 10^{-3}$ | $[1.5 \cdot 10^{-3}, 1.5 \cdot 10^{-3}]$ | $4.3 \cdot 10^{-3}$ | $[4.3 \cdot 10^{-3}, 4.4 \cdot 10^{-3}]$ | $\mu m^2 s^{-1}$ |
| $\omega_{de \rightarrow D,E}$ | $2.0 \cdot 10^{-1}$ | $[2.0 \cdot 10^{-1}, 2.0 \cdot 10^{-1}]$ | $2.9 \cdot 10^{-1}$ | $[2.8 \cdot 10^{-1}, 3.0 \cdot 10^{-1}]$ | $s^{-1}$ |
| $\omega_{de \rightarrow D,e}$ | $2.2 \cdot 10^{-4}$ | $[0, 5.4 \cdot 10^{-4}]$ | $2.7 \cdot 10^{-7}$ | $[0, 1.7 \cdot 10^{-2}]$ | $s^{-1}$ |
| $\omega_{de \rightarrow d,e}$ | $6.0 \cdot 10^{-2}$ | $[6.0 \cdot 10^{-2}, 6.0 \cdot 10^{-2}]$ | $5.9 \cdot 10^{-2}$ | $[0, 7.3 \cdot 10^{-2}]$ | $s^{-1}$ |
| $\omega_{ded,e \rightarrow de,de}$ | 9.9 | [9.9, 9.9] | $2.1 \cdot 10^{-14}$ | $[0, 2.0 \cdot 10^{-3}]$ | $\mu m^2 s^{-1}$ |
| $\omega_{ded \rightarrow d,de}$ | $2.5 \cdot 10^{-2}$ | $[2.4 \cdot 10^{-2}, 2.7 \cdot 10^{-2}]$ | 8.6 | [8.5, 8.6] | $s^{-1}$ |
| $\omega_{e \rightarrow E}$ | $5.6 \cdot 10^{-2}$ | $[5.6 \cdot 10^{-2}, 5.6 \cdot 10^{-2}]$ | $5.5 \cdot 10^{-2}$ | $[5.3 \cdot 10^{-2}, 5.6 \cdot 10^{-2}]$ | $s^{-1}$ |

**Table S6:** Parameters from the fits of the FHNM to the oscillation data and the MinD dissociation data.

| Parameter | Oscillation Data |  | MinD Dissociation Data |  | Units |
| --- | --- | --- | --- | --- | --- |
|  | Value | 95% Confidence Interval | Value | 95% Confidence Interval |  |
| $C_d$ | $1.0 \cdot 10^2$ | $[6.4, 1.9 \cdot 10^2]$ | $1.7 \cdot 10^2$ | $[0, 1.7 \cdot 10^2]$ | $\mu m^{-2}$ |
| $C_d$ (w/o MinE) | | | 0 | [0, 0] | $\mu m^{-2}$ |
| $C_e$ | $9.6 \cdot 10^1$ | $[6.2 \cdot 10^1, 1.3 \cdot 10^2]$ | 0 | $[0, 2.8 \cdot 10^1]$ | $\mu m^{-2}$ |
| $a$ | $1.4 \cdot 10^{-8}$ | $[1.1 \cdot 10^{-8}, 1.9 \cdot 10^{-8}]$ | $1.1 \cdot 10^{-11}$ | $[1.1 \cdot 10^{-11}, 1.3 \cdot 10^{-11}]$ | $\mu m^4 s^{-1}$ |
| $b$ | $2.0 \cdot 10^{-1}$ | $[1.7 \cdot 10^{-1}, 2.6 \cdot 10^{-1}]$ | $1.4 \cdot 10^{-1}$ | $[1.2 \cdot 10^{-1}, 1.5 \cdot 10^{-1}]$ | $s^{-1}$ |
| $c$ | $1.2 \cdot 10^{-2}$ | $[1.1 \cdot 10^{-2}, 1.2 \cdot 10^{-2}]$ | $1.6 \cdot 10^{-2}$ | $[1.5 \cdot 10^{-2}, 1.8 \cdot 10^{-2}]$ | $s^{-1}$ |
| $d$ | $1.4 \cdot 10^{-2}$ | $[1.3 \cdot 10^{-2}, 1.5 \cdot 10^{-2}]$ | $1.0 \cdot 10^{-2}$ | $[7.4 \cdot 10^{-3}, 1.4 \cdot 10^{-2}]$ | $s^{-1}$ |
| $I$ | $6.0 \cdot 10^1$ | $[4.6 \cdot 10^1, 7.8 \cdot 10^1]$ | 0 | [0, 3.3] | $\mu m^{-2} s^{-1}$ |
| $v_l$ | $9.7 \cdot 10^2$ | $[7.3 \cdot 10^2, 1.2 \cdot 10^3]$ | $1.3 \cdot 10^4$ | $[1.1 \cdot 10^4, 1.4 \cdot 10^4]$ | $\mu m^{-2}$ |
| $v_u$ | $7.5 \cdot 10^3$ | $[7.3 \cdot 10^3, 7.7 \cdot 10^3]$ | $7.8 \cdot 10^5$ | $[7.2 \cdot 10^5, 7.9 \cdot 10^5]$ | $\mu m^{-2}$ |

**Table S7:**
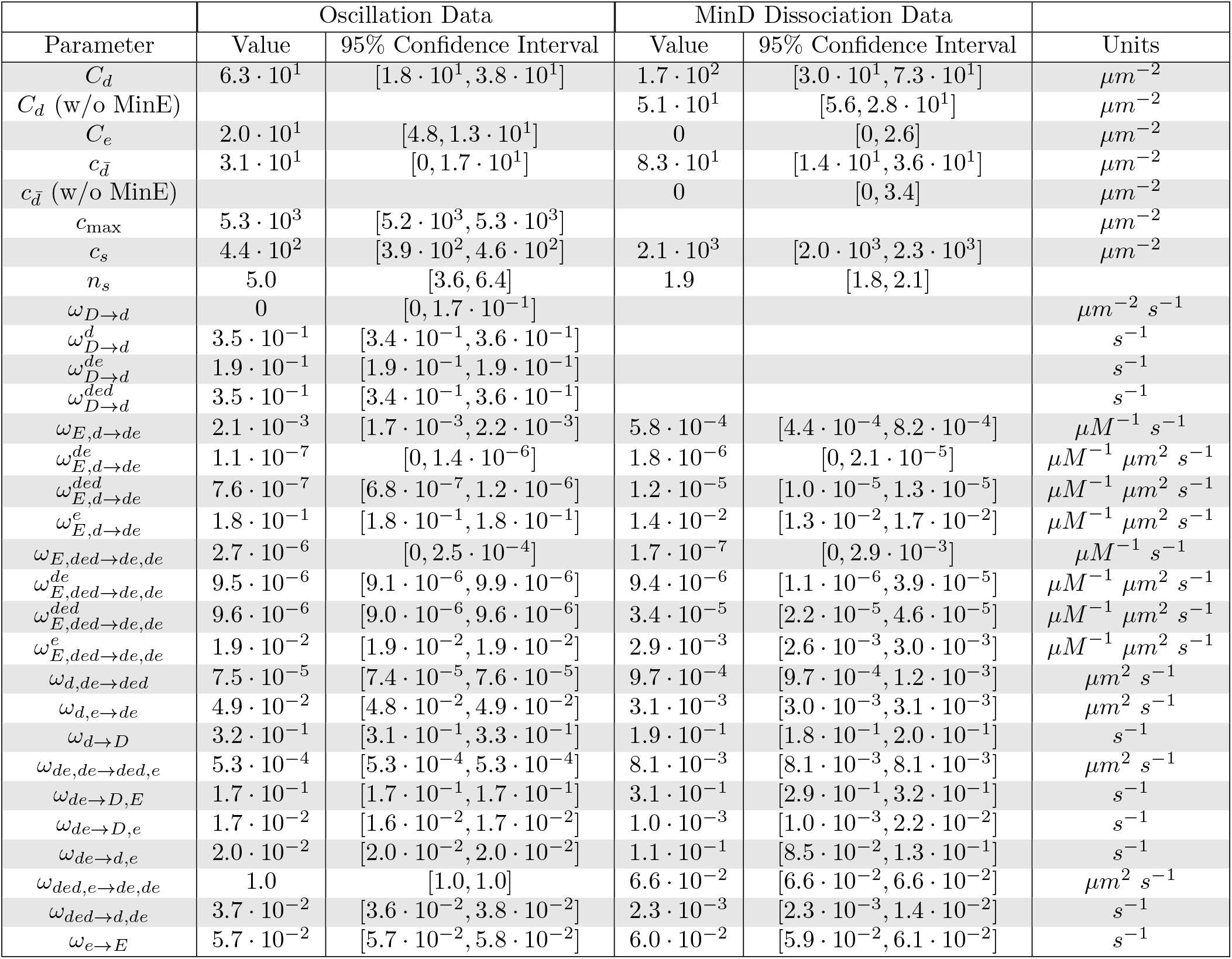
Parameters from the simultaneous fitting of the AABSM to the oscillation data and the MinD dissociation data with parameter constraints. *γ* = 16 in the constrained optimization. We presume that these are the most biologically relevant parameter estimates we have. See the caption of Fig. S9 for details of the constrained fit, notably regarding changes to the AABSM involving rate parameters 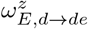 and 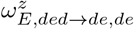 for *z* ∈ {*Ø, de, ded, e*}.

